# Conserved epigenetic regulatory logic infers genes governing cell identity

**DOI:** 10.1101/635516

**Authors:** Woo Jun Shim, Enakshi Sinniah, Jun Xu, Burcu Vitrinel, Michael Alexanian, Gaia Andreoletti, Sophie Shen, Yuliangzi Sun, Brad Balderson, Carles Boix, Guangdun Peng, Naihe Jing, Yuliang Wang, Manolis Kellis, Patrick P L Tam, Aaron Smith, Michael Piper, Lionel Christiaen, Quan Nguyen, Mikael Bodén, Nathan J. Palpant

## Abstract

Determining genes orchestrating cell differentiation in development and disease remains a fundamental goal of cell biology. This study establishes a genome-wide metric based on the gene-repressive tri-methylation of histone 3 lysine 27 (H3K27me3) across hundreds of diverse cell types to identify genetic regulators of cell differentiation. We introduce a computational method, TRIAGE, that uses discordance between gene-repressive tendency and expression to identify genetic drivers of cell identity. We apply TRIAGE to millions of genome-wide single-cell transcriptomes, diverse omics platforms, and eukaryotic cells and tissue types. Using a wide range of data, we validate TRIAGE’s performance for identifying cell-type specific regulatory factors across diverse species including human, mouse, boar, bird, fish, and tunicate. Using CRISPR gene editing, we use TRIAGE to experimentally validate *RNF220* as a regulator of *Ciona* cardiopharyngeal development and *SIX3* as required for differentiation of endoderm in human pluripotent stem cells. A record of this paper’s Transparent Peer Review process is included in the Supplemental Information.

## INTRODUCTION

Cellular identity is controlled by an interplay of regulatory molecules that cause changes in gene expression across the genome (Morris and Daley, 2013). Histone modifications (HMs) activate or repress genes to guide cell decisions during differentiation and homeostasis via mechanisms that are partially conserved across species (Alexanian et al., 2017; Boyer et al., 2006; Margueron and Reinberg, 2011; Nakamura et al., 2014). HMs have been found to be structurally and functionally linked to cell type specific genome architecture and gene regulation (Cahan et al., 2014; Rehimi et al., 2016). Histone 3 lysine 27 trimethylation (H3K27me3) is a chromatin mark deposited by the polycomb repressive complex-2 (PRC2) to suppress expression of genes (Margueron and Reinberg, 2011). The interplay of epigenomic control of gene expression by H3K27me3 and other activating histone marks, such as H3K4me3, guide cell lineage decisions to derive specific functional cell types (Van Handel et al., 2012). Computational methods using genome-wide measures of chromatin state and gene expression can therefore enable efficient prediction of genes controlling cell decisions (Benayoun et al., 2014; Rehimi et al., 2016; Whyte et al., 2013). These strategies have played critical roles in advancing fields of cell biology to inform the genetic basis of cell reprogramming and differentiation (Takahashi and Yamanaka, 2006).

Here, we demonstrate that a computational method formulated using the repressive tendency *via* H3K27me3 strongly predicts genes controlling cell differentiation decisions. The method draws on the principle that cell differentiation decisions are mediated in large part by selective epigenetic repression of regulatory genes (Stergachis et al., 2013). Genes that are repressed in many cell types are likely to play a key regulatory role for the rare cell types in which the gene is expressed. When measured across diverse cell types, the selective *absence* of broad H3K27me3 domains can therefore be used to predict cell type-specific genetic regulators. We show that the method can analyze millions of heterogeneous cell transcriptomes simultaneously to infer cell type-specific regulatory genes from diverse animal species. The approach we take departs from and complements analyses that require two or more relevant cellular conditions to be assayed. Instead, we draw on the cellular diversity represented in existing consortia epigenetic data as a background to evaluate genome-wide features of genetic regulation sourced from individual cell types. Analytical tools like this that evaluate patterns of genome regulation in the context of cellular diversity will enhance our capacity to understand the mechanistic basis of cellular heterogeneity in development, homeostasis and disease conditions.

## RESULTS

### Cell type specific regulatory genes have broad H3K27me3 domains in diverse cell types

We set out to test the hypothesis that analyzing the genomes of diverse cell and tissue types could be used to determine variation in epigenetic control of genes underpinning cell differentiation. We used NIH Epigenome Roadmap data (Kundaje et al., 2015), which contains ChIP-seq analysis of H3K4me3, H3K36me3, H3K27me3, H3K4me1, H3K27ac and H3K9me3 from 111 tissue or cell types **(Table S1)**. To associate HMs with genes, we linked the single broadest HM domain based on overlap with a RefSeq gene-body plus 2.5kb upstream of its transcriptional start site **(Figure S1A)**. For each HM, we found that the top 100 genes associated with the broadest domain were remarkably consistent between cell types **(Figure S1B)**, however, broad domains of different HMs marked distinct sets of genes **(Figure S1C)**.

We next assessed the breadth of histone domains in each of 111 samples as they correlate with genes that control cell type-specific functions. To establish a positive gene set for cell type-specific regulatory genes, we populated a list of 634 variably expressed transcription factors (VETFs) having a coefficient of variation greater than 1 across 46 NIH Epigenome Roadmap RNA-seq data sets (Perez-Lluch et al., 2015) (**Table S2**). We used Shannon entropy to quantify cell type specificity (Schug et al., 2005) and demonstrate that VETFs are significantly more cell type-specific, compared to non-VETFs (*p*=4.31e-230, Wilcoxon rank-sum test, one-tailed) or protein coding genes (*p*=1.55e-108) (**Figure 1A inset)** and their expression is more negatively correlated to the H3K27me3 breadth (**Figure S1D**). We showed that VETFs are highly consistent with a set of 713 tissue specific TFs independently identified from 233 tissue groups (*p<*2.2e-16, hypergeometric test) (D’Alessio et al., 2015). Taken together, VETFs provide a positive gene set where their enrichment is a performance metric for identifying cell type-specific regulatory genes.

**Figure 1:**
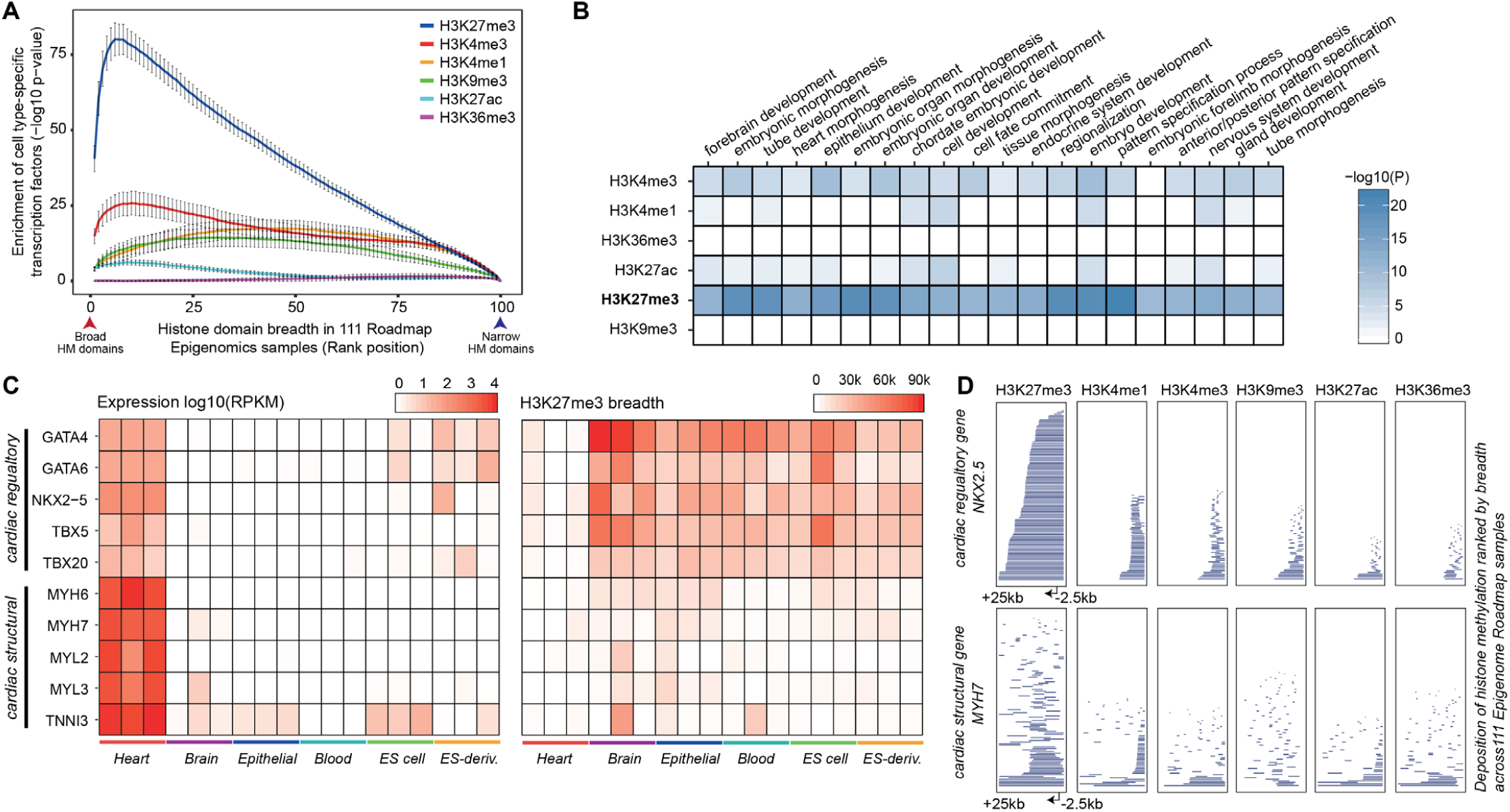
Tissue and cell type-regulatory genes are associated with broad H3K27me3. (A) Variably expressed transcription factors (VETFs) are strongly associated with broad H3K27me3 domains. Genes (*n*=26,833) are associated with a single broadest proximal histone modification (HM) domain and ranked by the breadth of the associated HM domain across 111 NIH Roadmap Epigenomics Project tissue or cell types. Genes are grouped into percentile bins. Mean enrichment of VETFs with the 95% confidence interval is shown (*y*-axis). *p*=6.66e-16 at top 5% broadest H3K27me3 domains by Fisher’s exact test, one-tailed. (B) Gene Ontology (GO) biological process enrichment of top 200 genes most frequently associated with top 5% broadest HM domains across 111 cell types (Fisher’s exact test, one-tailed). H3K27me3 broad domains are enriched for cell regulatory genes. (C) Gene expression level (left) and the H3K27me3 domain breadth (right) for selected cardiac-specific regulatory genes and structural genes across 18 Roadmap tissue samples; Heart (E095, E104, E105), Brain (E070, E071, E082), Epithelial (E057, E058, E059), Blood (E037, E038, E047), ES cell (E003, E016, E024) and ES-derived (E004, E005, E006). Gene expression distinguishes cardiac-specific genes whereas the selective absence of H3K27me3 broad domains distinguishes cardiac-specific regulatory genes from structural genes. (D) HM depositions in a cardiac regulatory gene *NKX2-5* and structural gene *MYH7* (−2.5kb upstream of the TSS to +25kb downstream). Data are ranked based on the size of histone domain with 111 Epigenome Roadmap cell types stacked vertically. Consistent deposition of broad H3K27me3 domains across diverse cell types demarcates cell type-specific regulatory genes from structural genes.

To calculate the enrichment of cell type-specific regulatory genes as they relate to HM domains, all HM domains assigned to genes were ranked by breadth and analyzed using Fisher’s exact test to assess enrichment of VETFs.

These data show that broad H3K27me3 domains are strongly enriched for VETFs **(Figures 1A, S1F)**, consistent with a correlation between repression of VETFs and breadth of H3K27me3 domains **(Figure S1E)**. Further supporting this, unsupervised analysis of genes marked by the broadest H3K27me3 domains are uniquely enriched in morphogenic and developmental regulators **(Figure 1B)**. These regulatory genes have broad H3K27me3 repression in many cell types to preclude them from being mis-expressed and therefore providing an epigenetic control mechanism controlling their expression. Taken together, this demonstrates that H3K27me3 broad domains as measured across diverse cell and tissue types provides a strategy to enrich for cell type-specific regulatory genes.

To illustrate the distinctive enrichment of H3K27me3 in regulatory genes as opposed to structural or housekeeping genes, we extracted expression and chromatin data from cardiomyocytes **(Figure 1C-D)**. The transcript abundance of cardiac regulatory transcription factors (i.e. *GATA4, GATA6, NKX2-5, TBX5* and *TBX20*) and structural sarcomere genes (i.e. *MYH6, MYH7, MYL2, MYL3* and *TNNI3*) are all elevated in cardiac cells compared to other cell types, but cannot be distinguished as regulatory or structural genes except by differential expression **(Figure 1C)**. In contrast, in all cell types *except* the heart, H3K27me3 domains broader than 30kb consistently identify cardiac regulatory genes from structural genes **(Figure 1C)**. No other HM analyzed demarcates cell type-specific regulatory genes from structural genes in this manner **(Figures 1D, S1G)**. This establishes the rationale that genes having cell type-specific regulatory functions can be identified based on the frequency of H3K27me3 across the locus in heterogeneous cell types.

### The repressive tendency of a gene defines a genome-wide metric predictive of cell type-specific regulatory genes

We established a simple, quantitative logic that leverages the significance of broad H3K27me3 domains for distinguishing regulatory genes: Broad H3K27me3 domains occurs predominantly over critical cell type-specific regulatory genes; depositing this mark sets the default gene activity to “off” such that cell type-specific expression requires selective removal of H3K27me3 (Boyer et al., 2006; Lee et al., 2006). Conversely, genes with housekeeping or non-regulatory roles rarely host broad H3K27me3 domains.

To implement this concept, we calculated for each gene in the genome across 111 NIH epigenome cell and tissue types (i) the total lengths (breadths) of H3K27me3 domains in base-pairs and multiplied this by (ii) the proportion of cell types in which the gene’s H3K27me3 breadth is within the top 5% of broad domains **(Figure 2A)**; this is justified on the basis of the 4∼8% thresholds that show similar, high level of recovery of regulatory factors (**Figures 1A and S2A)**. This analysis assigns a single value to every gene we call its repressive tendency score (RTS) that defines its association with broad H3K27me3 domains **(Table S3)**. Using the NIH Epigenome Roadmap data, the RTS is calculated for 99.3% (or 26,833 genes) of all RefSeq genes. To demonstrate that this formulation is agnostic to the composition of cell types, we note that for all genes, the RTS is within one standard deviation of the mean of bootstrapping empirical distribution derived from 10,000 re-samplings of cell types. We note that the 111 cell types provide sufficient sample size to calculate a stable RTS, independent of the peak calling method to define H3K27me3 domains (**Figure S2B-C**), with over 85% of assigned H3K27me3 domains overlapping only a single protein-coding gene **(Figure S2D)**. Using a subsampling approach, we show that the RTS can be calculated from approximately as few as 60 randomly selected but distinct cell and tissue types (**Figures S2F-G**). The gene-centric measure is an aggregate of broad H3K27me3 domains around each gene across diverse cell types. Making use of the recently released EpiMap data comprising 18 genome-wide features from 834 cell and tissue types (Adsera et al., 2019), we show that RTS is highly consistent regardless of whether the input samples are comprised of cell lines, embryonic or adult samples, or healthy and diseased cell types **(Figure S2H)**.

**Figure 2:**
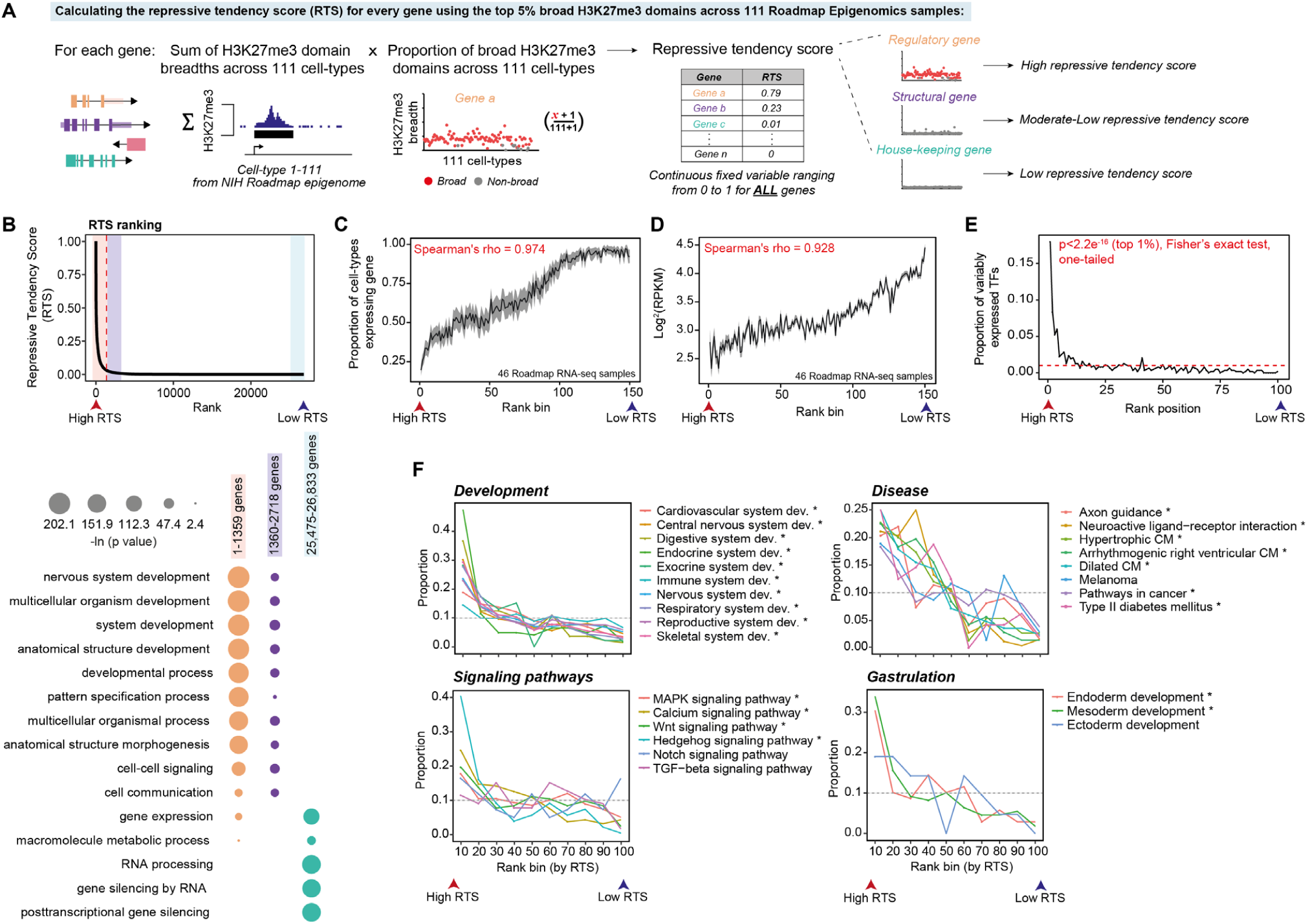
Genes governing development are frequently associated with broad H3K27me3 domains. (A) Schematic diagram showing steps for using 111 NIH Roadmap Epigenomics samples to calculate the repressive tendency score (RTS) for protein coding genes. (B) (Top) Distribution of the RTS. Red dashed line is the inflection point on the interpolated curve (RTS=0.03) above which genes exhibit increased RTS (*n*=1,359). (Bottom) GO biological process enrichment in genes ranked by the RTS (Fisher’s exact test, one-tailed) (Table S4 for the full list). (C) High RTS ranked genes are associated with cell/tissue specificity. Each rank bin includes 100 genes and used to calculate the proportion of cell types where a given gene is expressed (RPKM>1). The average proportion is calculated for each rank bin with the 95% confidence interval shown. (D) High RTS genes tend to be lowly expressed. Each rank bin includes 100 genes and the average expression value for each bin is shown with the 95% confidence interval. (E) Variably expressed transcription factors (VETFs) are significantly associated with a high RTS (for example, *p*<2.2e-16 at the first rank position (top 1%), Fisher’s exact test, one-tailed). Each rank bin includes 1% of all RefSeq genes with RTS (*n*=26,833). Red dashed line shows the uniform distribution (proportion=0.01). (F) High RTS genes are enriched in regulators of development and disease processes across all organ systems (Table S4 for the full list). Each rank bin includes 1% of all RefSeq genes (*n*=26,833). Asterisks indicate significant enrichment of a given GO term within the top 10% genes (Benjamini-Hochberg FDR<0.05, Fisher’s exact test, one-tailed).

Using RTS values above the inflection point (RTS>0.03) of the interpolated RTS curve, we identified a priority set of 1,359 genes. These genes show significant enrichment for cell type-specific, lowly expressed regulators of cellular diversification including organ development, pattern specification and multicellular organismal processes **(Figure 2B-D)**. Among the 1,359 priority genes, we identified enrichment of VETFs (odds ratio=13.85, *p=*3.2e-151, Fisher’s exact test, one-tailed), homeobox proteins (Zhong and Holland, 2011) (odds ratio=37.42, *p=*7.98e-135) and KEGG signaling genes (Kanehisa and Goto, 2000) (odds ratio=2.66, *p*=1.73e-13) (**Figure 2E)**. The priority set also comprises non-coding RNAs including known regulators of development such as FENDRR and HOTAIR (Grote and Herrmann, 2013; Rinn et al., 2007). Furthermore, genes with high RTS values are enriched in regulators of biological processes including gastrulation and organ morphogenesis, and comprise members of major signaling pathways, and genetic determinants of diverse pathologies including cardiovascular disease, diabetes, neurological disorders and cancer **(Figure 2F, Table S4)**. Additionally, we evaluated the biological significance of perturbing genes with a high or low RTS by analyzing an independent data-resource containing 714 transgenes (including 481 TFs) conditionally over-expressed in hESCs subsequently analyzed by RNA-seq or microarray (**Figure S3A**) (Nakatake et al., 2020). Overexpression of TRIAGE priority genes resulted in a significantly higher number of differentially expressed genes, showing greater perturbation of the transcriptome, compared to non-priority genes (*p*≤0.0001, two-tailed) **(Figures S3B-C)**. This suggests that genes with high RTS are regulators of cell identity. Ranking genes based on RTS is a simple strategy to enrich for genetic factors controlling cell type-specific differentiation and function.

### Applying the RTS to orthologous gene expression data infers regulatory genes of that cell

The transcriptome of a cell is comprised of diverse cell type-specific structural, housekeeping and regulatory genes. The expression of regulatory genes, such as transcription factors (TFs), is difficult to detect, and changes are harder to quantify due to their typical low abundance, relative to structural and housekeeping genes. To enable genes to be ranked based on their regulatory potential, we sought to calculate an adjusted abundance measure for a gene that accounts for the gene’s expression level observed in a foreground population of cells or single cell, and its tendency of being epigenetically repressed across a background of cell types. For any gene *i* the product between its expression value (*Y*_*i*_) and repressive tendency score (*R*_*i*_) gives rise to its discordance score (*D*_*i*_). This defines a method we call TRIAGE (Transcriptional Regulatory Inference Analysis of Gene Expression):

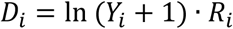

The discordance score (DS) represents a gene’s tendency to be epigenetically repressed and the observed transcriptional abundance of that gene in orthologous input gene expression data. TRIAGE does not require an index cell type reference; instead it uses the repressive tendency collected from a diverse spectrum of cell and tissue types, to linearly amplify or attenuate the measure of gene abundance from *any* orthologous input sample **(Figures 3A, S2E, S3D-E)**.

**Figure 3:**
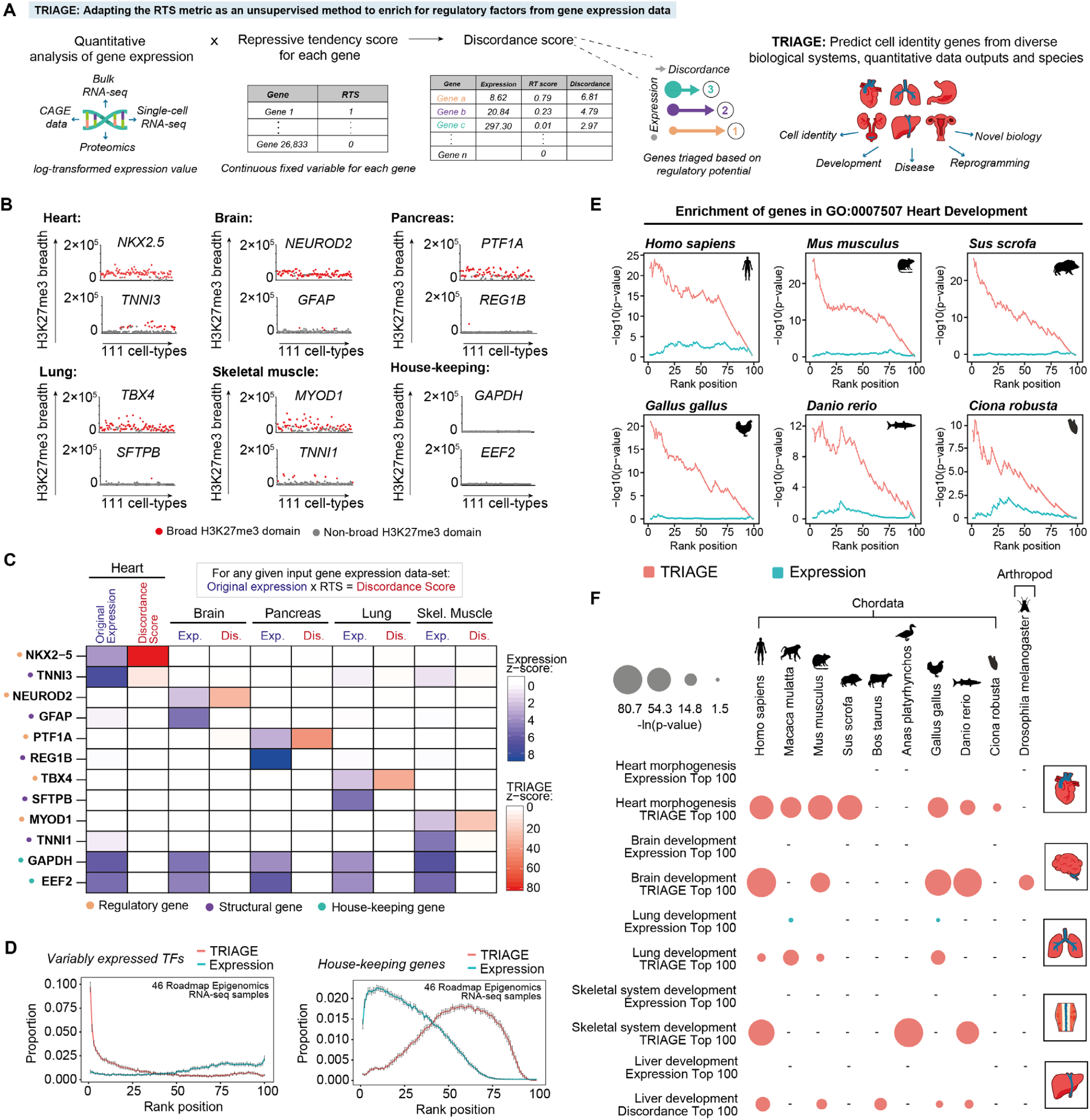
TRIAGE: an unsupervised strategy to infer cell type-specific regulatory factors from gene expression input alone. (A) Schematic illustrating TRIAGE analysis of gene expression input data to generate a discordance score (DS) for prioritization of genes based on regulatory potential from orthologous gene expression data set. (B) Regulatory genes with broad H3K27me3 domains (in base-pairs) as exemplified by selected tissue type-specific regulatory, structural, and housekeeping genes. (C) TRIAGE prioritizes tissue type-specific regulatory genes and reduces the relative abundance of housekeeping and structural genes. Average expression values (*Exp*.) from selected GTEx tissue samples transformed by TRIAGE (*Dis*.). (D) Across 46 Roadmap Epigenomics samples, TRIAGE significantly enriches for VETFs (*p*<2.2e-16, Fisher’s exact test, one-tailed) but reduces the proportion of housekeeping genes (*p*<2.2e-16, Wilcoxon rank-sum test, one-tailed) at top 1% RTS rank position. Average values from Roadmap cell types are shown, with the 95% confidence interval scale bar. (E) TRIAGE enriches for regulators of heart development (GO:0007507) among genes ranked by the TRIAGE discordance score (DS) (red) compared to original expression (cyan) in cardiac RNA-seq samples from diverse animal species (Fisher’s exact test, one-tailed). (F) TRIAGE enriches for tissue-specific developmental GO terms among top 100 genes ranked by the DS (red) compared to original expression (cyan) across diverse animal species (Fisher’s exact test, one-sided). Hyphen (-) indicates no data set available.

As expected, analysis of cell-specific genes across 111 NIH Epigenome Roadmap samples shows that H3K27me3 broad domains reproducibly mark regulatory genes as opposed to structural or housekeeping genes **(Figure 3B)**. Using RNA-seq data transformed with RTS, we show that TRIAGE efficiently reduces the relative abundance of structural and housekeeping genes, while enriching for regulatory genes in a cell type-specific manner **(Figures 3C, S3F)**. TRIAGE transformation of 46 Roadmap RNA-seq samples results in enrichment of tissue or cell type-specific TFs among the top 1% in every cell type while it reduces the relative abundance of housekeeping genes **(Figure 3D)**. Analysis of the Pearson correlation distances between Roadmap tissue types (Scornavacca et al., 2011) shows that TRIAGE increases the similarity between samples from the same tissue by ∼29% when compared to distances calculated using absolute expression levels **(Figure S3G)**.

### RTS identifies cell identity genes from diverse tissues and species

Genetic mechanisms that control cell decisions are highly evolutionarily conserved. Using inter-species gene mapping, we tested whether TRIAGE could identify regulatory drivers of heart development across diverse chordate species including mammals (i.e. *Homo sapiens, Mus musculus*, and *Sus scrofa*), bird (*Gallus gallus*), fish (*Danio rerio*) and in vertebrate tunicate (*Ciona robusta*) **(Figure 3E)**. In contrast to expression alone, TRIAGE accurately recovered cardiac regulatory genes across all species. More broadly, we used TRIAGE to enrich for relevant tissue morphogenesis biological processes from diverse cell types and species including arthropods **(Figure 3F)**. While TRIAGE is currently devised using human epigenetic data, these data show that TRIAGE can be used to identify regulatory genes from cell types that are conserved across the animal kingdom.

### RTS analyses infer regulatory control points of disease

Cell differentiation decisions in development are commonly re-activated in disease contexts to drive cell differentiation decisions in response to cell stress (Rajabi et al., 2007). Indeed, analysis of high-RTS genes shows enrichment in disease-related KEGG pathways and ClinVar disease terms, including both congenital and acquired disorders **(Figure S4A)**. Next, we used TRIAGE to analyze two disease data sets: single-cell RNA-seq data of melanoma (Tirosh et al., 2016), and RNA-seq data of hearts where pre-established heart failure (transverse aortic constriction, TAC) was treated with JQ1, a small molecule BET inhibitor known to prevent pathological cardiac remodeling (Anand et al., 2013; Duan et al., 2017). We show that among the top ranked genes, TRIAGE consistently prioritizes genes with known involvement in skin development and melanoma pathogenesis using independently derived positive gene sets (Tirosh et al., 2016; Verfaillie et al., 2015) **(Figure S4B-C)**, as well as enrichment of stress-associated gene ontology pathways **(Figure S4D-E)**. TRIAGE-based ranked genes highlighted the potent anti-fibrotic effect of JQ1 without the use of a canonical differential expression analysis **(Figure S4E)**. While disease responses are complex and involve denovo chromatin changes not represented in the reference healthy cell and tissue types collated in the Epigenome Roadmap, these data demonstrate that TRIAGE can provide a strategy to study mechanistic processes in disease.

### TRIAGE provides a unique vantage point into genomic data

We compared the data analysis pipelines of TRIAGE with various benchmark genomic analysis methods that also rely on reference to epigenetic information including functional heterogeneity (FH) analysis (Rehimi et al., 2016), H3K4me3 broad domains (Benayoun et al., 2014) and super-enhancers (Jiang et al., 2019; Whyte et al., 2013) **(Figure 4A)**. TRIAGE is unique among these analysis approaches by requiring only gene expression data without the need for epigenetic sequencing input. Notably, comparison of the overlap of genes prioritized by these approaches across different tissue types reveals largely non-overlapping genes sets (**Figure 4B, Table S5**). For example, among genes identified within each method, TRIAGE distinctively detects the largest number of cardiac transcription factors, many known to be central regulators of cardiac morphogenesis including *IRX4, GATA5, TBX5, MSX1*, and *EOMES* (**Figure 4B**) (Russ et al., 2000; Takeuchi and Bruneau, 2009; Waardenberg et al., 2014). Consistent with this, TRIAGE performs well when compared with these methods in terms of sensitivity and precision at identifying cell type-specific regulatory genes **(Figures 4C-D, S5A-D, Tables S6-8)**. These data demonstrate that TRIAGE captures gene sets distinct to other epigenetic analysis strategies and therefore provides an effective and complementary approach for evaluating genomic data in coordination with routine analysis pipelines.

**Figure 4:**
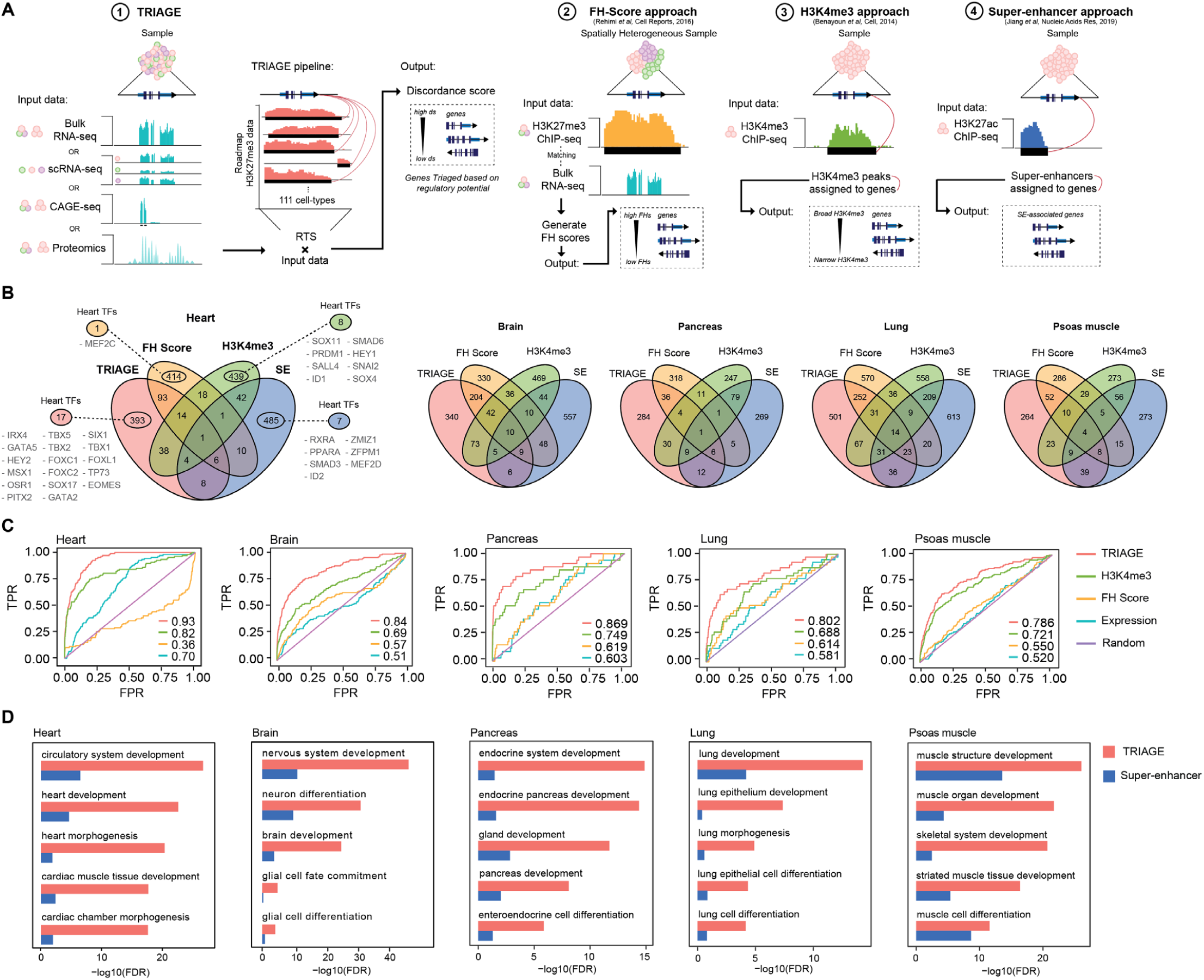
TRIAGE identifies unique genomic regulatory features that complement those of other methods. (A) Schematic diagrams of computational approaches to infer cell identity genes using epigenomic and/or transcriptomic data with TRIAGE, FH Score, H3K4me3, or super-enhancers. (B) Venn diagrams demonstrate that different epigenetic prediction methods capture distinct gene sets across multiple tissues. (C) Receiver operating characteristic (ROC) plots indicate that TRIAGE consistently recovers tissue-specific regulatory genes. Area under curve (AUC) values are shown on the right bottom corner of the plot. (D) TRIAGE improves on super-enhancers prioritizing genes for developmental and morphogenesis process across diverse cell and tissue types.

### RTS reveals the mechanistic basis of cell heterogeneity at single-cell resolution

Recent developments in barcoding and multiplexing have enabled scalable analysis of thousands to millions of cells (Cao et al., 2019). Determining mechanistic information from diverse cell subtypes captured using single-cell analytics remains a challenge. TRIAGE does not require epigenetic data for a cell of interest, making it applicable to any bulk or single-cell transcriptomic data input.

We analyzed 43,168 cells captured across a 30 day time-course of *in vitro* cardiac-directed differentiation from human pluripotent stem cells (hPSCs) (Friedman et al., 2018). Analysis of day-30 cardiomyocytes using standard expression data show high abundance genes dominated by housekeeping and sarcomere genes, whereas TRIAGE efficiently identifies cardiomyocyte regulatory genes including *NKX2-5, HAND1, GATA4, IRX4* **(Figure 5A-B)**. Notably, TRIAGE retains highly expressed cell type-specific structural genes providing an integrated readout of genes involved in cell regulation and function **(Figure 5C)**. We used TRIAGE to simultaneously convert single-cell expression data comprising ten different cell subpopulations spanning gastrulation stage, progenitor and definitive cell types **(Figure 5D)**. In contrast to expression data, which significantly enriches for structural and housekeeping genes, TRIAGE consistently identifies gene sets associated with developmental regulation of diverse and biologically distinct subpopulation through differentiation **(Figures 5E, S6A-B)**. We show that differential expression analysis results in outcomes that depend heavily on the comparison cell type, whereas TRIAGE identifies population-specific regulatory genes without external reference comparisons **(Figure 5F)**. Lastly, we show that TRIAGE predictions are not explained merely by prioritizing expressed TFs **(Figure S5E-F, Table S9)**, indicating the capability of TRIAGE to filter out housekeeping TFs that do not govern cell type-specific functions.

**Figure 5:**
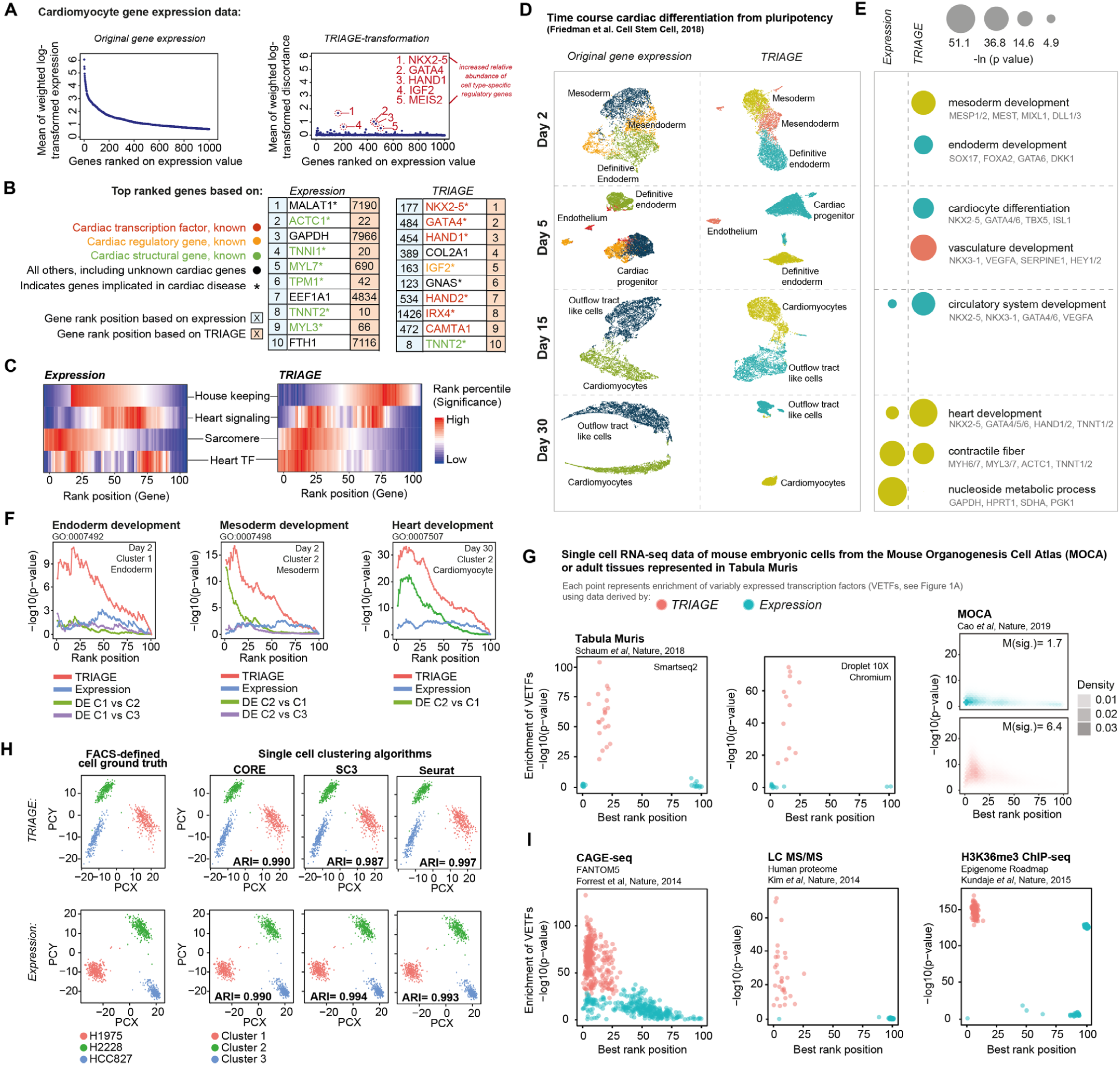
TRIAGE enables analyses of large single-cell and multi-omics data sets. (A) RNA-seq analysis of cardiomyocytes based on RNA expression (left) and gene discordance scores after TRIAGE analysis (right) where genes are ordered by their expression. TRIAGE reveals known cardiac regulators that are lowly ranked based on original input gene expression. (B) Top 10 genes ranked based on their RNA expression (left) and TRIAGE (right) analyses of hiPSC derived cardiomyocytes (Friedman et al., 2018). (C) Distribution of selected groups of genes, including housekeeping, heart signaling (genes with the heart development term GO:0007507 and at least one KEGG signaling pathway term), sarcomere (genes with the sarcomere term GO:0030017) and heart TFs (TFs with the heart development term GO:0007507) when ranked by their RNA expression value (left) and gene discordance score after TRIAGE analysis (right). Each rank bin includes 1% of all expressed genes in the data set. Enrichment of a given group is calculated for each rank bin relative to all genes (Fisher’s exact test, one-tailed). (D) Analyses of single-cell RNA-seq over a time course of cardiac differentiation from pluripotency by clustering of cells from RNA expression (left) and TRIAGE (right). (E) During *in vitro* mesendoderm differentiation (*y*-axis) RNA expression enriches for structural and housekeeping genes, and TRIAGE enriches for Gene Ontology’s Biological Process terms related to cell type-specific regulatory developmental processes (Fisher’s exact test, one-tailed). (F) TRIAGE (red) enriches for developmental terms among top ranked genes compared to expression (blue) or differential gene expression (green or purple). (G) TRIAGE enriches for VETFs across diverse scRNA-seq sequencing platforms (i.e. Smart-seq2 or Droplet 10X chromium) and scales to analyze millions of cells in the mouse organogenesis cell atlas (MOCA). Genes are sorted by either RNA expression or TRIAGE, and then grouped into a percentile bin. Enrichment of VETFs for each sample is summarized by the most significant *p*-value (*y*-axis) at the corresponding rank bin position (*x*-axis). (H) TRIAGE accurately clusters scRNA-seq data compared to original RNA expression data evaluated using ground truth analysis captured using Mixology data sets (Tian et al., 2019). (I) TRIAGE used for analysis of diverse data types and separation of VETFs where gene expression is quantified, including CAGE-seq, proteomics, and H3K36me3 tag density.

### TRIAGE is applicable to diverse omics data types and scales to large single-cell data sets

We tested the utility of TRIAGE using different genomic and proteomic data types. Using the Tabula Muris data of nearly 100,000 cells from 20 different mouse tissues at single-cell resolution (Schaum et al., 2018), TRIAGE consistently enriches for cell type-specific regulatory genes compared to original expression with no difference between droplet and smartseq2 data sets **(Figure 5G, Table S10)**. Analysis of the mouse organogenesis cell atlas (MOCA (Cao et al., 2019)) data demonstrates that TRIAGE prioritizes cell type-specific regulatory genes in a scalable manner across more than 1.3 million mouse single-cell transcriptomes (*p*<2.2e-16, Wilcoxon rank-sum test for both median significance and rank, one-tailed, **Figure 5H)**. Lastly, we evaluated single-cell clustering accuracy based on ground truth (Tian et al., 2019) to assess the performance of TRIAGE using three independent algorithms (i.e. CORE, sc3, and Seurat). We show no difference in accurately assigning cells to the reference (ARI > 0.98) using original expression or TRIAGE transformed expression **(Figure 5I)**.

We hypothesized that TRIAGE could be used to study any genome-wide quantitative measurement of gene expression. We applied TRIAGE to 17,382 GTEx (v8) samples covering diverse cell and tissue types (Lonsdale et al., 2013). Compared to original expression values, TRIAGE prioritizes genes with tissue specific developmental functions (**Figure S7A-B, Table S11**). This finding is consistently observed across diverse omics data types that measure gene abundance. For example, TRIAGE outperforms original abundance metrics when measuring chromatin methylation for H3K36me3, a surrogate of RNA polymerase II activity deposited across gene bodies (Barski et al., 2007) collected from the 111 Roadmap samples **(Table S12, Figure 5J)**. We used data from FANTOM5 (D’Alessio et al., 2015; Forrest et al., 2014) to analyze cap analysis of gene expression (CAGE), a measure of genome-wide 5’ transcription activity. These data showed TRIAGE enriches for tissue-specific TFs (**Figures 5J, S8A-C, Table S13**) as well as tissue-specific GO biological process terms compared to CAGE input data (**Table S14**). Lastly, analysis of a draft map of the human proteome shows that TRIAGE enriches for regulatory drivers of 30 different tissue types from high resolution Fourier transform mass spectrometry data (Kim et al., 2014) **(Table S15)**. These findings illustrate that TRIAGE predicts regulatory drivers of cells using diverse genome-wide multi-omic endpoints.

### TRIAGE for novel gene discovery

We set out to demonstrate TRIAGE as an engine for gene discovery. We used a multi-step strategy involving analysis of TRIAGE-derived discordance score, RTS, and prior literature to identify candidate genes governing cell differentiation **(Figure S9A)**. We focused on transcription factors and signaling molecules which are known to act as upstream regulators of broad gene networks guiding cell differentiation decisions. Using data from single-cell analysis of cardiac differentiation (Friedman et al., 2018) we analyzed day 2 subpopulations and identified diverse, known regulatory genes governing mesendoderm cell differentiation **(Figure 6A)**. Among the TRIAGE identified genes was *SIX3*, a member of the sine oculis homeobox transcription factor family (RTS=0.54) **(Figures 6A-B, S9A-B)**. Though the r ole of *SIX3* in neuroectoderm specification has been studied extensively, little is known about its role in other germ layer derivatives (Carl et al., 2002; Lagutin et al., 2003; Steinmetz et al., 2010). Analysis of *SIX3* in hPSC *in vitro* cardiac differentiation shows robust expression in day 2 mesendoderm cell populations **(Figures 6C, S9C-D**). We analyzed the spatiotemporal transcriptional data from germ layer cells of mouse embryos (Peng et al., 2016), spanning pre-gastrula stages (E5.5) to late-gastrulation (E7.5) mouse embryos (**Figure S9E**). Spatio-temporal expression of *SIX3* is observed in the epiblast and neuroectoderm (Carl et al., 2002; Lagutin et al., 2003; Steinmetz et al., 2010) as well as early endoderm lineages (**Figures 6D, S9F)**. We also identified *SIX3* as a gene detected in a study of 434 individuals with brain development abnormalities (holoprosencephaly) with 8% of individuals found to have congenital heart disease. A patient with deletion of *SIX3* was diagnosed with dextrocardia (Tekendo-Ngongang et al., 2020), a condition where the heart develops properly except its position is reversed in the body further suggesting a potential direct or indirect role for *SIX3* in governing critical mesendoderm cell differentiation decisions.

**Figure 6:**
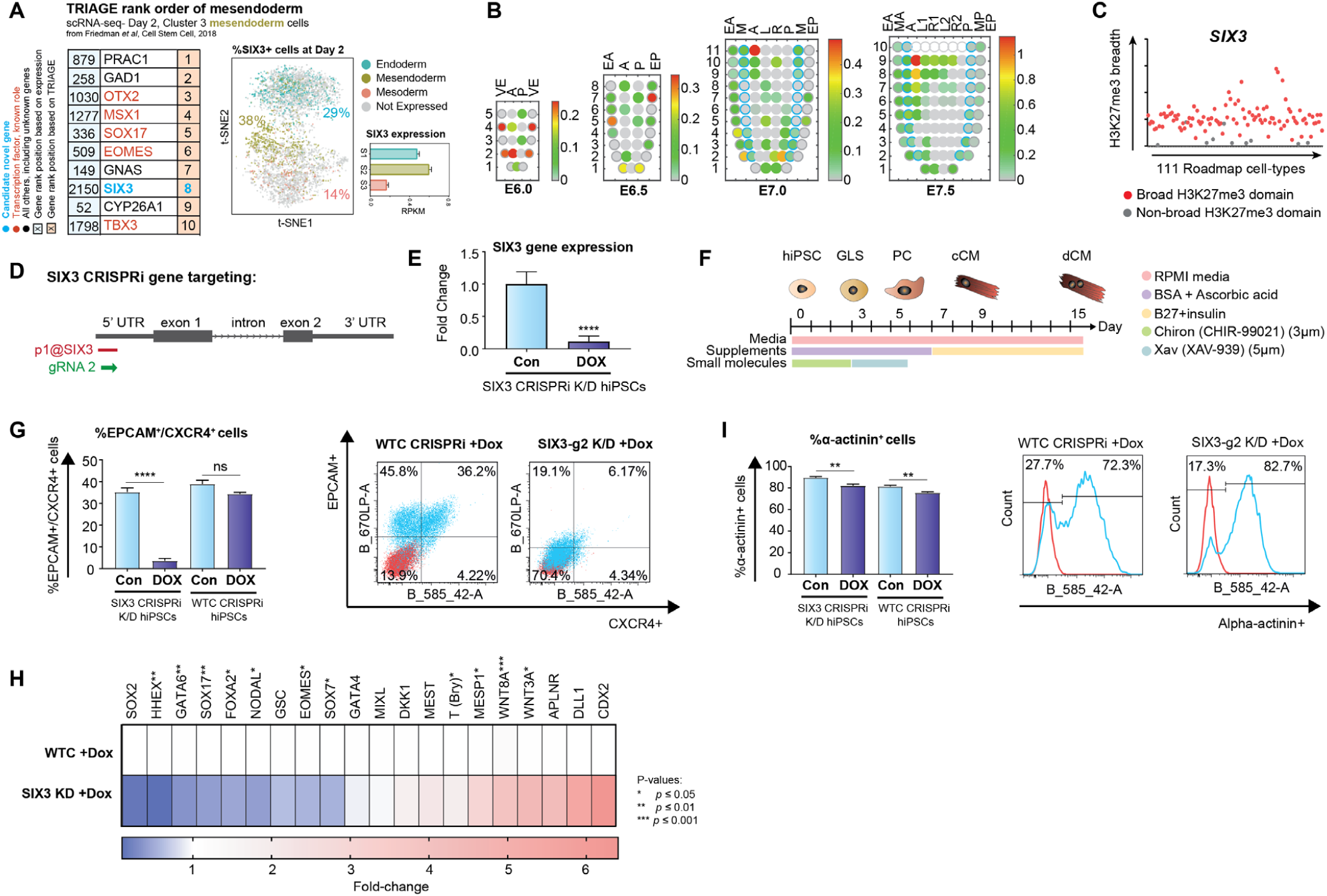
TRIAGE as an engine for gene discovery. (A) Top 10 genes ranked by RNA expression (left) and TRIAGE (right) from day-2 *in vitro* hPSC-derived mesendoderm, highlighting *SIX3* as a candidate gene identified by TRIAGE. t-SNE plot shows percentage of cells expressing *SIX3* and gene expression level of *SIX3* in different single-cell derived subpopulations. Genes labeled as “novel” have not been implicated in this process to the best of our knowledge. (B) Corn plots showing the spatial domain of *SIX3* expression in the germ layers of E5.5-E7.5 mouse embryos. Positions of the cell populations (‘‘kernels’’ in the 2D plot of RNA-seq data) in the embryo: the proximal-distal location in descending numerical order (1 = most distal site) and in the transverse plane of the germ layers: endoderm, anterior half (EA) and posterior half (EP); mesoderm, anterior half (MA) and posterior half (MP); epiblast/ectoderm, anterior (A), posterior (P) containing the primitive streak, right (R)-anterior (R1) and posterior (R2), left (L) – anterior (L1) and posterior (L2). (C) Breadths of H3K27me3 domains (in base-pairs) associated with *SIX3* gene across 111 Epigenome Roadmap samples. (D) Schematic overview of *SIX3* gene targeting by CRISPRi for conditional knockdown (KD) showing the position of gRNAs blocking CAGE-defined TSS of *SIX3*. (E) qPCR analysis of *SIX3* transcript abundance in control (con) vs *SIX3* CRISPRi KD iPSCs (DOX) (*n*=6-14 technical replicates per condition from 3-6 experiments). (F) Schematic of hiPSC directed *in vitro* cardiac differentiation protocol. (G) Day-2 FACS analysis of endoderm markers EPCAM/CXCR4 between control and dox-treated conditions in *SIX3* CRISPRi KD iPSCs and WTC GCaMP CRISPRi iPSCs are shown (*n*=12-16 technical replicates per condition from 4-5 experiments). *SIX3* CRISPRi KD iPSCs show significant (*p*<0.001) reduction in EPCAM^+^/CXCR4^+^ cells compared to dox-treated control iPSCs (WTC GCaMP CRISPRi). (H) qPCR analysis showing significant decreases in endoderm and mesendoderm markers and increases in mesoderm markers, respectively, in *SIX3* CRISPRi KD iPSCs compared to control (*n*=6-14 technical replicates per condition from 3-6 experiments). (I) Analysis of cardiomyocytes by FACs for α-actinin at day 15 of *in vitro* differentiation. Changes in α-actinin^+^ cells between control and dox-treated conditions in *SIX3* CRISPRi KD iPSCs and WTC GCaMP CRISPRi iPSCs are shown (*n*=6 technical replicates per condition from 3 experiments). *SIX3* CRISPRi KD iPSCs show no change in α-actinin^+^ cells compared to dox-treated control iPSCs (WTC GCaMP CRISPRi).

**Figure 7:**
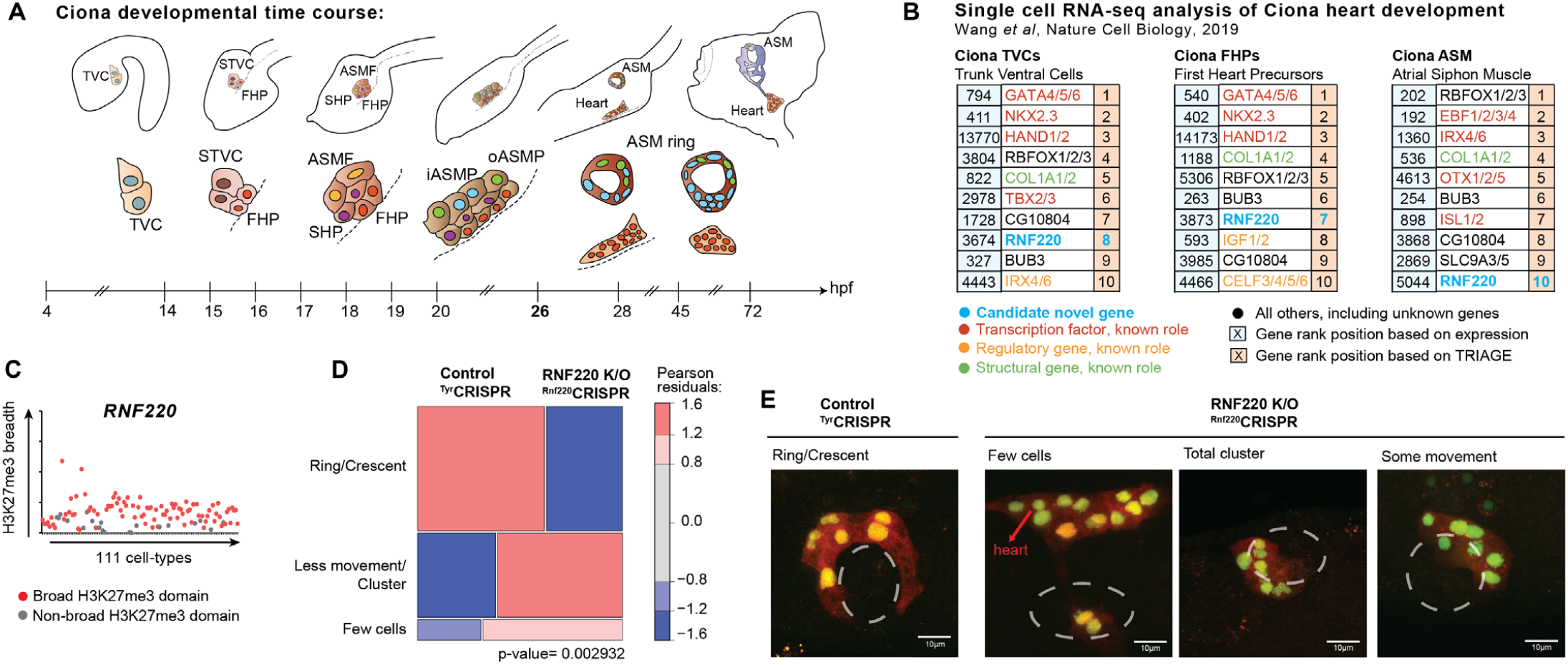
TRIAGE identifies regulators of heart development in the tunicate *Ciona*. (A) Schematic overview of cardiac development in *Ciona* from 4 to 72 hours post fertilization (hpf) at 18°C. Adapted from (Evans Anderson and Christiaen, 2016). TVC: trunk ventral cells; STVC: second TVC; FHP: first heart precursor; SHP: second heart precursor; ASMF: atrial siphon muscle founder cells; iASMP: inner atrial siphon muscle precursor; oASMP: outer atrial siphon muscle precursor. (B) Top 10 genes ranked by expression value (left) or TRIAGE (right) from populations found during *Ciona* heart development *in vivo*, highlighting *RNF220* as a candidate gene. Genes labeled as “novel” have not been implicated in this process to the best of our knowledge. (C) Breadths of H3K27me3 domains (in base-pairs) associated with *RNF220* gene across the 111 NIH Epigenomes data sets. (D-E) Mosaic plots (D) and images (E) showing ASM precursor phenotypes at 26 hpf labeled with Mesp>H2B:GFP and Mesp>mCherry in control knockout and *RNF220*-knockout animals (*n*=100). *p*-value represents the chi-sq test between two experimental conditions. Images in (E) derived from *Ciona robusta* cardiopharyngeal mesoderm.

To test this hypothesis, we established CRISPRi loss-of-function hPSCs in which *SIX3* transcription is blocked at its CAGE-defined transcription start site (TSS) in a dox-dependent manner **(Figures 6E-F)**. Cells were differentiated and analyzed at day 2 **(Figure 6G)**. We found that *SIX3* loss-of-function depleted endoderm and pan-mesendoderm genes (*n*=6-14 technical replicates per condition from 3-6 experiments) with FACS data indicating specific depletion of CXCR4^+^/EPCAM^+^ endoderm cells (*n*=12-16 technical replicates per condition from 4-5 experiments) **(Figures 6H-K, S9G, S10)**. In contrast, genes marking various mesodermal lineages (pan-mesoderm, paraxial mesoderm and lateral mesoderm) were either not affected or upregulated upon loss of *SIX3* **(Figure 6H)**. Consistent with this, FACS analysis of alpha-actinin^+^ cardiomyocytes on day 15 of differentiation showed no difference between *SIX3*-knockdown cells compared to dox-treated controls (*n*=6 technical replicates per condition from 3 experiments) **(Figures 6L-N, S9H)**. Taken together, this finding provides the first evidence linking *SIX3* to mesendoderm cell differentiation and situates this gene into mechanisms of cardiac malformation that are consistent with genetic defects in endoderm that are known to cause heart patterning defects (Viotti et al., 2012).

Lastly, we used TRIAGE to identify previously unknown developmental regulators in a distant chordate species, *Ciona robusta*. Single cell RNA-seq data comprising cell subpopulations captured across a time-course of cardiac development were analyzed with TRIAGE using a customized gene mapping tool to link human to *Ciona* genes **(Figure 6O)** (Wang et al., 2019). The top ranked genes based on TRIAGE were analyzed **(Figures 6P, S9A)**. *RNF220* (RTS=0.30, **Figure 6Q**), an E3 ubiquitin ligase governing Wnt signaling pathway activity through β-catenin degradation (Ma et al., 2014), was identified as a regulatory gene not previously implicated in cardiopharyngeal development. Utilizing CRISPR control vs. *RNF220*-knockout, we demonstrate that *Mesp* lineage progenitors of control animals form the expected ring of pharyngeal muscle progenitors around the atrial siphon placode, whereas *RNF220*-knockout embryos showed morphogenetic defects (*n*=100) **(Figures 6R-S)**. Collectively, these data illustrate that TRIAGE identifies functional regulatory determinants; we demonstrate how TRIAGE can support discovery of previously unknown biology and mechanisms of development.

## DISCUSSION

TRIAGE builds on the prevailing view that the epigenome guides cell regulatory processes. The selective absence of broad H3K27me3 domains at promoters is first established from diverse cell types and epigenetic information hosted by NIH Roadmap Epigenomics (Kundaje et al., 2015) and represented by a simple gene-specific repressive tendency score. The RTS demarcates 5% of Refseq genes that are consistently regulated by broad H3K27me3 domains across major human cell lineages; we show this priority set is highly enriched in genes that govern cell differentiation and lineage diversification.

Secondly, TRIAGE exploits the discordance between any gene’s expression and its RTS to predict cell-specific genes regulating cell identity with sensitivity and precision. We demonstrate a capacity of TRIAGE to prioritize genes involved in cell differentiation decisions in development and disease contexts. These inferences establish a genetic basis for gaining mechanistic insights into cell decisions, including the discovery of previously unknown genetic regulators as demonstrated by gene loss of function studies.

The scalability and versatility of TRIAGE is exemplified in its implementation across diverse data types including single cell RNA-seq, ATAC-seq, CAGE-seq, proteomics, ChIP-seq or any genomic analysis method linked to protein coding genes. We provide evidence for TRIAGE as a tool for studying the genetic basis of cell differentiation choices in cell types from every organ system in the body and across species. TRIAGE provides a unique vantage point for studying genomic data that is complementary to many routine analysis approaches such as differential expression. While we demonstrate its use for studying developmental and cell differentiation decisions, additional studies are merited for evaluating the role of TRIAGE in analysis of adult tissues and disease responses.

The evolutionary conservation of epigenetic regulation suggests that the repressive tendency can be applied across eukaryotic cell types where gene expression is governed by the polycomb group complex-2. PRC2 and its regulation of histone methylation are known to govern genes in protists, animals, plants as well as fungi (Margueron and Reinberg, 2011). Indeed, zebrafish (Wu et al., 2011) and medaka (Nakamura et al., 2014) genes with broad H3K27me3 deposition at promoter sites encode master developmental regulators overlapping with those found in our study. This illustrates the conservation of PRC2-mediated H3K27me3 regulation and repression of genomic loci across species and its role controlling cell identity (Boyer et al., 2006; Fujikura et al., 2002).

In contrast to previous strategies (Cahan et al., 2014; Rackham et al., 2016), TRIAGE does not require comparison against external reference information akin to differential gene expression analysis. It detects known regulatory genes that are distinct from genes identified using other approaches such as FH Score (Rehimi et al., 2016), H3K4me3 broad domains (Benayoun et al., 2014), or super-enhancers (Creyghton et al., 2010; Hnisz et al., 2013; Jiang et al., 2019; Whyte et al., 2013). TRIAGE provides a seamless interface with a range of input data types measuring gene outputs of a cell with application by genetic orthologs for the study of any cell type across the animal kingdom.

Among the limitations of TRIAGE, we used RTS mapped specifically to protein coding genes. This limits TRIAGE inference predictions to the 2% of the genome encoding proteins and precludes analysis of intergenic domains that also encode regulatory information (Andersson et al., 2014). Second, we note limitations on cross-species applications based on gene mapping tools and the profound differences in composition of cellular diversity that distinguish animal phyla. Despite this limitation, we demonstrate cross-species predictions and functional evidence of gene discovery in distantly related chordates using RTS for human. Third, TRIAGE uses a single number assigned to each protein coding gene to prioritise genes from cell type-specific data and therefore is not designed to evaluate changes between samples as would occur with fold change measured by differential expression. Finally, while the biological diversity represented in the 111 NIH Roadmap Epigenomics is limited, we illustrate that the method is robust to the sample input, by performing analyses on epigenomic data from >830 cell types in EpiMap. We anticipate that the generation of new reference epigenome databases (Adsera et al., 2019) covering a broader range of cell and tissue-types will refine and expand the use of RTS revealing stress-sensitive loci and novel disease drivers.

This study establishes a unique strategy for the study of the transcriptional control mechanisms governing cell diversification, without demanding the provision of multiple data sets. With expanded analysis of epigenomic information of diverse cell and tissue types across eukaryotic species, the analysis can be implemented to study mechanisms underlying the genetic basis of complex cell traits, as well as gaining insights into evolutionary biology and genetic adaption across eukaryotic genomes governed by PRC2.

## METHODS

### Data sets

We compared associations of 6 different histone modifications (H3K4me1, H3K4me3, H3K9me3, H3K27me3, H3K27ac and H3K36me3) with genes using consolidated broad peak representations from the NIH Roadmap database (Kundaje et al., 2015). We extracted a curated set of 1,605 human TFs from Animal TFDB and DBD (Wilson et al., 2008; Zhang et al., 2015). We identified 3,804 housekeeping genes from the published literature (Eisenberg and Levanon, 2013). The repressive tendency score of genes were estimated based on biological diversity represented in the 111 NIH Roadmap human tissue and cell types.

We utilized publicly available expression datasets to test performance of TRIAGE. We applied TRIAGE to 17,382 transcriptome samples from GTEx (v8) (Lonsdale et al., 2013), Roadmap and 329 selected FANTOM CAGE-seq developmental samples (**Table S1**) (Forrest et al., 2014). We demonstrated multi-omics applicability of TRIAGE with single-cell transcriptomes (Cao et al., 2019; Friedman et al., 2018; Schaum et al., 2018), human proteome (Kim et al., 2014) and Roadmap H3K36me3 ChIP-seq data.

We evaluated TRIAGE in prioritizing cell identity genes against approaches based on H3K4me3 peak breadth (Benayoun et al., 2014), functional heterogeneity (FH) score (Rehimi et al., 2016), super-enhancers (SEs) (Hnisz et al., 2013; Jiang et al., 2019; Whyte et al., 2013) and differentially expressed (DE) gene analysis. To this end, we selected 5 distinct tissue-groups as the reference point; (i) Brain, (ii) Lung, (iii) Pancreas, (iv) Skeletal muscle and (v) Heart. We calculated metrics using transcriptomic abundance and ChIP-seq data from the Roadmap data; Brain germinal matrix (E070) for the brain, Lung (E096) for the lung, Pancreas (E098) for the pancreas, Psoas muscle (E100) for the skeletal muscle and a published data set for cardiac progenitor cells for the heart (GSE97080) (Palpant et al., 2017b). We collated nearest active genes associated with SEs in corresponding tissue-groups (See **Performance analysis of TRIAGE against existing methods**) (Jiang et al., 2019). We also used GTEx expression profiles of samples that belong to these 5 tissue-groups; 417 samples from the left ventricle (Heart), 195 samples from the cortex (Brain), 268 samples from the pancreas (Pancreas), 607 samples from the lung (Lung) and 718 samples from the skeletal muscle (Skeletal muscle). We averaged gene expression values of samples for each tissue-group.

To identify a gene set regulating the cell fate, we used gene ontology (GO) annotation data (Ashburner et al., 2000). We identified TFs with a tissue-specific GO biological process (BP) term as the positive gene set with a defined regulatory role. For instance, we used ‘heart development’ term (GO:0007507) to select a set of TFs specific for heart development. Similarly, ‘brain development’ (GO:0007420), ‘lung development’ (GO:0030324), ‘pancreas development’ (GO:0031016) and ‘muscle structure development’ (GO:0061061) GO terms were used to collect regulatory gene sets for brain, lung, pancreas and skeletal muscle samples, respectively. Finally, we extracted expressed genes (i.e. RPKM>1) in a given input sample as an active set of positive genes.

### Identifying genes as a proxy for cell type-specific regulatory genes

Along with TFs with a tissue-type-specific GO BP term, we identified a gene set that universally represents cell type-specific regulatory genes. We identified 634 TFs whose expression values shows substantial variation (i.e. coefficient of variation>1) across the 46 Roadmap cell types and labelled them as variably expressed TFs (VETFs) (**Table S2**). This classification is based on previous observations that expression of developmentally regulated genes is highly variable both temporally and spatially (Perez-Lluch et al., 2015). Collectively, these 634 VETFs encompass regulatory genes for a broad range of cell and tissue types. We used a subset of these TFs that were expressed (RPKM>1 or equivalent) in a given input sample as the positive gene set.

To ensure that our analysis is not confined to the narrow definition of cell type-specific regulatory genes above, we collated curated sets of tissue type-specific TFs (D’Alessio et al., 2015). This study ranked human TFs for their tissue specificity across 233 tissue groups based on the expressional specificity. We took top the 20 TFs for each tissue group by their specificity score, yielding a total of 713 tissue-specific TFs. 428 of 634 our VETFs were indeed identified as members of these TFs, demonstrating agreement with the VETFs (*p<*2.2e-16, hypergeometric test). We used these tissue-specific TF sets as a complementary source of the positive gene set to assess performance of TRIAGE with the CAGE expression data.

### Quantifying cell type specificity of VETFs

We use Shannon entropy to quantify the specificity of expression for VETFs, as observed across 46 Roadmap cell types (Schug et al., 2005). The *relative* expression is calculated as

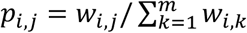

where *w*_*i,j*_ is the expression value of gene *i* in cell type *j*, from 46 Roadmap cell types (*m*=46) and its cell type specificity is

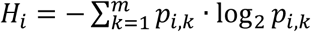

The cell type specificity ranges between 0 (when the gene is uniquely expressed in a single cell type) and log_2_ *m* (when the gene is expressed uniformly across all cell types). VETFs had significantly lower entropies (mean=3.64), compared to non-VETFs (mean=5.25, *p*=4.31e-230, Wilcoxon rank-sum test, one-tailed) and all protein-coding genes (mean=4.48, *p*=1.55e-108), indicating their expressional specificity.

### Association of genes with a histone modification (HM) domain

To assign peaks (referred to as domains hereafter due to the focus on broad peaks) to genes, we used following steps.

1. Defining the proximal region of the gene. We defined a proximal region for each gene. The proximal region for a gene is the RefSeq gene body plus a region extended by 2.5kb from the TSS in the upstream direction.
2. Provisional assignment of domains to genes. For each gene, we first identified HM domains with their center position overlapping the proximal region. These domains were provisionally assigned to the corresponding gene. Domains that were broad, with their center position outside of the proximal region were still included if the domain overlapped with any proximal regions of genes, in which case, the domain was provisionally assigned to all overlapping genes. Suppose that gene *i* in cell type *j* have a set of provisionally assigned domains, *D*_*i,j*_ = {*d*_1_, *d*_2_, …, *d*_*l*_, … } where *d*_*l*_ is the breadth (in base-pairs) of the *l-*th domain provisionally assigned to the gene.
3. Final assignment of domains to genes. If multiple domains were assigned to a gene *i* in cell type *j*, it is represented by the breadth of the broadest domain

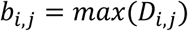

This yields a single gene assigned with a single domain in a given cell type. We used BEDTOOLS intersect program to assign HM domains to RefSeq genes (Quinlan and Hall, 2010). We removed any intergenic domains that did not overlap any proximal regions of genes to reduce potential bias from including intergenic regulatory elements (Rada-Iglesias et al., 2011). We use a term *assigned* to indicate that the HM domain has been linked to the gene.

Our approach to annotate genes with breadth of the broadest H3K27me3 domain did not result in loss of relevant functional information. To illustrate this point, we calculated correlation between expression level of a gene and (i) the breadth of the broadest H3K27me3 domain or (ii) the sum of breadths of all H3K27me3 domains assigned to the gene. When we performed this analysis across all the cell types in Roadmap, we found no significant difference in the correlation between these two approaches, with the mean Spearman’s *rho* of −0.364 and −0.337 respectively. Furthermore, the majority of domains finally assigned to protein-coding genes (i.e. approximately 85% of 1,537,514 assigned H3K27me3 domains across the 111 cell types) overlapped a single gene (**Figure S2C**). Approximately 66.9% of all H3K27me3 domains assigned to genes were identified in the RefSeq gene region while remaining 22.6% and 10.5% were in the promoter (+2.5kb from the TSS) and intergenic regions respectively, indicating that the majority (89.5%) of the assigned H3K27me3 domains were proximal to the gene.

For cell type *j*, we have a set of HM domain breadth values *B*_*j*_ = {*b*_1,*j*_, *b*_2,*j*_, …, *b*_*i,j*_, …, *b*_*n,j*_} for *n* genes. Subsequently, we normalized breadth values to yield the breadth score (*h*) for all genes across the cell types, 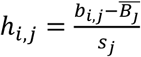 where 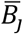 and *s*_*j*_ are the sample mean and the standard deviation of HM domain breadths in cell type *j*.

### Genomic locations of H3K27me3 domain

We investigated genomic locations of assigned 1,537,514 H3K27me3 domains used for the RTS calculation. First, we identified a center position of each domain and used BEDTOOLS intersect program to identify if they overlap with (i) known RefSeq genes or (ii) promoters (defined as +2.5kb from the RefSeq TSS). If the domain does not overlap any of these, we labelled the domain as (iii) intergenic.

### Defining the broad H3K27me3 domain

We defined *broad* domains as the top 5% broadest domains, assigned to any gene. This threshold was based on efficacy of the threshold to recover genes with known regulatory and signaling functions. First, we observed that VETFs were strongly associated with broad H3K27me3 domains across the Roadmap cell types. When genes were ranked by the breadth of their finally assigned H3K27me3 domain, VETFs were most significantly (*p*=6.66e-16, Fisher’s exact test, one-tailed) enriched in the top 5% across all cell types (**Figure 1C**). To decide on this threshold, we assessed sensitivity of different thresholds (or rank position) to recover the VETFs and 641 KEGG signaling genes. We calculated a detection ratio (i.e. number of domains assigned to any positive gene divided by the total number of domains drawn at a given threshold) and a recovery percent (i.e. number of positive genes identified by collecting domains from all 111 Roadmap cell types at a given threshold, divided by the total number of positive genes). As each gene is assigned with a single domain in a given cell type, the detection ratio ranges from 0 (i.e. no single domain is associated with the positive gene) to 1 (i.e. all domains are associated with the positive gene). Similarly, the recovery percentage represents a cumulative proportion of positive genes captured at a threshold. We sought a threshold that maximized the detection ratio while recovering a majority of positive genes. Our analysis showed that a threshold at the 95th percentile (or top 5% broadest) met such conditions for both gene sets; only minor variations were observed at nearby rank positions (**Figure S2A**). For instance, approximately 81% of VETFs are identified at the 95th percentile (or rank position 5) where more than 15% of domains are assigned to VETFs. A variable (*X*_*i,j*_) represents a binary outcome of whether a gene *i* is assigned with a H3K27me3 domain that is in the top 5% broadest in a cell type *j*.

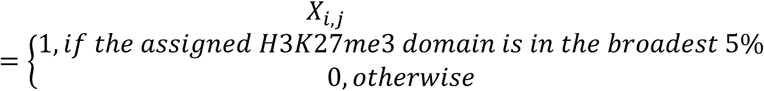

### Association of genes with broad HM domains

To understand association of broad HM domains with genes, we ranked genes by breadth of the assigned domain and grouped them into bins of 100 genes across the cell types. Jaccard similarity index was calculated at each rank bin between all pairs of cell types. To identify genes that are frequently associated with assigned broad HM domains across the Roadmap cell types, we counted the number of cell types where a given gene was assigned with the broad HM. We ranked genes by the count and identified the top 200 genes for each HM type.

### Estimating H3K27me3 repressive tendency of the gene

We hypothesized that the regulatory importance of a gene in any cell type can be determined from evidence of (i) expression level of that gene in the same cell type, and (ii) breadth of H3K27me3 domains collected from a diverse range of cell types. To test this, we proposed a method that first quantifies the association of a gene with the H3K27me3 domain. For each gene, the method considers (i) a sum of H3K27me3 breadth scores for the gene calculated from *m* cell types (e.g. 111 Roadmap cell types) and (ii) the number of cell types where the gene is associated with a broad H3K27me3 domain. For gene *i*, sum of the breadth scores (*v* _*i*_) is defined as follows.

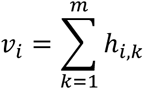

The sum of breadth scores was then re-scaled into a range from 0 to 1 (*v*_i_ ′) as follows, where *v*_max_ and *v*_min_ are maximum and minimum sums of the breadth scores from all genes respectively.

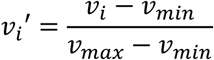

The association score of gene *i* with the H3K27me3 domain (*a*_*i*_) was then calculated as the product of the scaled sum of breadth scores (*v*_*i*_ ′) and a proportion of cell types where the gene was associated with the broad H3K27me3 domain. To include genes without any broad H3K27me3 domains in any cell types, we added a pseudo-count of 1.

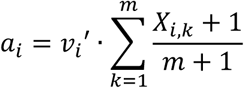

Finally, we re-scaled the association score into a range of 0 (min) and 1 (max) to obtain the repressive tendency score (RTS) for the gene. For gene *i*, RTS is defined as follows, where *m*_*max*_ and *m*_*min*_ are maximum and minimum association scores for all genes, respectively.

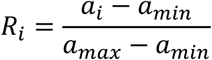

### Consistency of the RTS values between different data sources

To understand how the biological composition of samples affects calculation of the RTS, we utilised EpiMap (Adsera et al., 2019). We downloaded H3K27me3 data for all 834 EpiMap samples in the BigWig format. These were converted to bedgraph format using UCSC bigWigToBedgraph program, obtained from http://hgdownload.soe.ucsc.edu/admin/exe/. To identify H3K27me3-enriched regions, we used MACS2 peak calling program, with -bdgbroadcall option to capture broad deposition of H3K27me3 (Zhang et al., 2008). We identified H3K27me3-enriched regions across all 834 EpiMap samples.

We calculated RTS values using (i) all 834 EpiMap samples, (ii) 332 embryonic samples (i.e. ‘embryonic’ in life stage based on the EpiMap metadata annotation), (iii) 389 adult samples (i.e. ‘adult’ in life stage), (iv) 122 cancer samples (i.e. ‘cancer’ in group), or (v) 216 cell line samples (i.e. ‘cell line’ in type).

### Performance analysis of TRIAGE against existing methods

We extensively analyzed performance of TRIAGE against existing computational methods using various metrics. These include metrics based on (i) breadth of H3K4me3 peaks proximal to the gene (Benayoun et al., 2014), (ii) breadth of H3K27me3 peaks at the gene locus multiplied by the corresponding expression level in a spatially heterogenous cell population (Rehimi et al., 2016) and (iii) super-enhancer (SE) (Hnisz et al., 2013; Jiang et al., 2019; Whyte et al., 2013) as well as common practices of differentially expressed gene (DEG) analysis.

Furthermore, we tested TRIAGE with publicly available datasets including GTEx transcriptomes (v8) (Lonsdale et al., 2013), FANTOM5 CAGE-seq data (Forrest et al., 2014) as well as single-cell transcriptomes (Friedman et al., 2018; Schaum et al., 2018), human proteomes (Kim et al., 2014) and Roadmap H3K36me3 ChIP-seq data sets (Kundaje et al., 2015) encompassing diverse cell and tissue groups to compare the performance of TRIAGE against the gene expression readouts (See **Benchmarking TRIAGE in multi-omics platforms**). The following analyses were performed to performance test TRIAGE against previous epigenetic prediction analysis strategies:

#### a. H3K4me3 breadth

To compare performance of TRIAGE against the H3K4me3 breadth-based metric (Benayoun et al., 2014), we used the 5 distinct tissue groups (See **Datasets**). H3K4me3 peaks were assigned to nearest RefSeq genes using an in-house Python script. Peaks located further than 2.5kb from any RefSeq TSSs were excluded. For genes with multiple assigned peaks, the broadest H3K4me3 peak was used to annotate the gene. Subsequently, genes were ranked by the breadth of the H3K4me3 peak.

#### b. Functional heterogeneity (FH) score

To test performance of TRIAGE on predicting developmental regulators against FH score (Rehimi et al., 2016), we used chicken embryo dataset for enriched H3K27me3 domains identified in Rehimi’s study and the gene expression data (GSE89606) (Rehimi et al., 2016). These datasets satisfy the assumption of spatially heterogenous gene expression to calculate the FH score. We also downloaded genome annotation file (GFF3 format) for galGal4 chicken genome assembly (https://asia.ensembl.org/info/data/ftp/index.html) to quantify breadth of H3K27me3 peaks at the gene using BEDTOOLS intersect program (Quinlan and Hall, 2010). Furthermore, while the assumption of spatially heterogeneous gene expression is not strictly met, we also included FH score to evaluate its performance and applicability on the selected 5 distinct tissue groups (See Datasets).

Genes were ranked at both developmental time-points (HH14 and HH19) based on 3 metrics; (i) normalized gene expression value (TPM), (ii) FH score from Rehimi’s study, and (iii) DS from TRIAGE. A total of 13,214 genes with an expression value (TPM>0) were included. We analyzed enrichment of 19 related GO BP terms used in Rehimi’s study using Fisher’s exact test (one-tailed) **(Figure S5B)**.

#### c. Super-enhancer (SE) based approach

We downloaded lists of SEs for 5 selected tissue-types from a published SE database SEdb (http://www.licpathway.net/sedb/index.php) (Jiang et al., 2019); (i) Heart left ventricle, (ii) Lung_30y, (iii) Pancreas, (iv) Psoas muscle_30y, and (v) Brain astrocyte which were linked to Roadmap epigenomes E095 (Left ventricle), E096 (Lung), E098 (Pancreas), E100 (Psoas muscle) and ENCODE astrocyte (Brain) respectively. The algorithm to define SEs, ROSE (Rank-Order of Super-Enhancers) gives a binary outcome (i.e. SE or not) for each enhancer element (Whyte et al., 2013). As such, we extracted all nearest active genes of SEs and compared their functional enrichment against the same number of highly ranked genes by TRIAGE (Left ventricle (*n*=557), Lung (*n*=955), Pancreas (*n*=382), Psoas muscle (*n*=409) and Brain (*n*=689)) using Fisher’s exact test (one-tailed).

#### d. Differentially expressed gene (DEG) analysis

To test performance of TRIAGE against DEG analysis, we used published single-cell transcriptomes for *in vitro* cardiac-directed differentiation (Friedman et al., 2018) as well as selected bulk RNA-seq data for 3 distinct tissue groups (i.e. Blood, Brain and Heart) from the Roadmap project (Kundaje et al., 2015).

We obtained cardiac single-cell transcriptomes for differentiation days 2 and 30 from the ArrayExpress database (E-MTAB-6268). The data were processed and cells were clustered as previously described (Friedman et al., 2018). For the Roadmap data, we extracted RNA-seq data for 3 representative samples for each tissue group (Blood; E037, E038, E047, Brain; E070, E071, E082 and Heart; E095, E104, E105). We calculated mean expression values of genes within each group or cell cluster. Genes were ranked by the fold change (FC) of the expression value between different groups or clusters using R library DESeq2 (Love et al., 2014). We also tested performance of TRIAGE with the input gene set restricted to only TFs. To this end, we first excluded all non-TF genes from the input data. While DEG analysis often focuses on differentially expressed TFs as a candidate regulatory gene set, TRIAGE offers an unsupervised approach to prioritize cell type-specific regulatory genes, without requiring any prior knowledge on the gene set.

To demonstrate ability of TRIAGE as a complementary method to DE analysis to identify causal factors of cell identity, we compared rank analysis of genes using 108 FANTOM5 CAGE-seq expression samples (as read counts) from 12 cell (or tissue) types **(Table S1)**. We used a TSS with the highest read count as an expression value for the gene. We performed the DE analysis for all possible pairs of these 12 cell types using DESeq2 R package (Love et al., 2014). Briefly, for each cell type, we identified top 100 significantly highly expressed genes by comparing to each of all other cell types (i.e. 12 * 11 = 132 pair-wise comparisons).

We first extracted genes that were significantly highly expressed (Benjamini-Hochberg FDR < 1e-06) in a given cell type compared to the other, then ranked those genes by the fold change. We used top 20 tissue specific TFs as the positive gene set for each cell (or tissue) type. We counted positive hits for each comparison and calculated the enrichment using hypergeometric test. For the comparison, we ran TRIAGE independently for the same set of the 12 cell types. Top 100 genes were identified by the discordance score and overlaps with the positive gene set were computed.

#### e. H3K27me3 gain/loss function

We tested how TRIAGE performs against a simple gain/loss H3K27me3 function using a published dataset for induced *in vitro* differentiation of human cardiovascular cells between two different time-points; before differentiation (day0) and definitive cardiovascular cell stage (day14) (Paige et al., 2012). To this end, we first obtained AffyExon microarray expression (GSE19090) and H3K27me3 ChIP-seq (GSE35583) data. We averaged probeset values to obtain the gene expression value and merged H3K27me3 peaks from replicates. To quantify H3K27me3 depositions, we mapped peaks to regions spanning from 2.5kb upstream of RefSeq TSSs to the entire gene body, which is represented by the number of overlapped base-pairs. We normalized depositions by the size of the region to get a value ranging between 0 (complete absence of H3K27me3) and 1 (completely covered by H3K27me3). Finally, change in the H3K27me3 deposition between the two time-points was calculated for all genes (i.e. Δ*H*3*K*27*me*3 = *H*3*K*27*me*3_*day*14_ − *H*3*K*27*me*3_*day*0_). We ranked genes by loss of the H3K27me3 signal, DS and the expression values at day14.

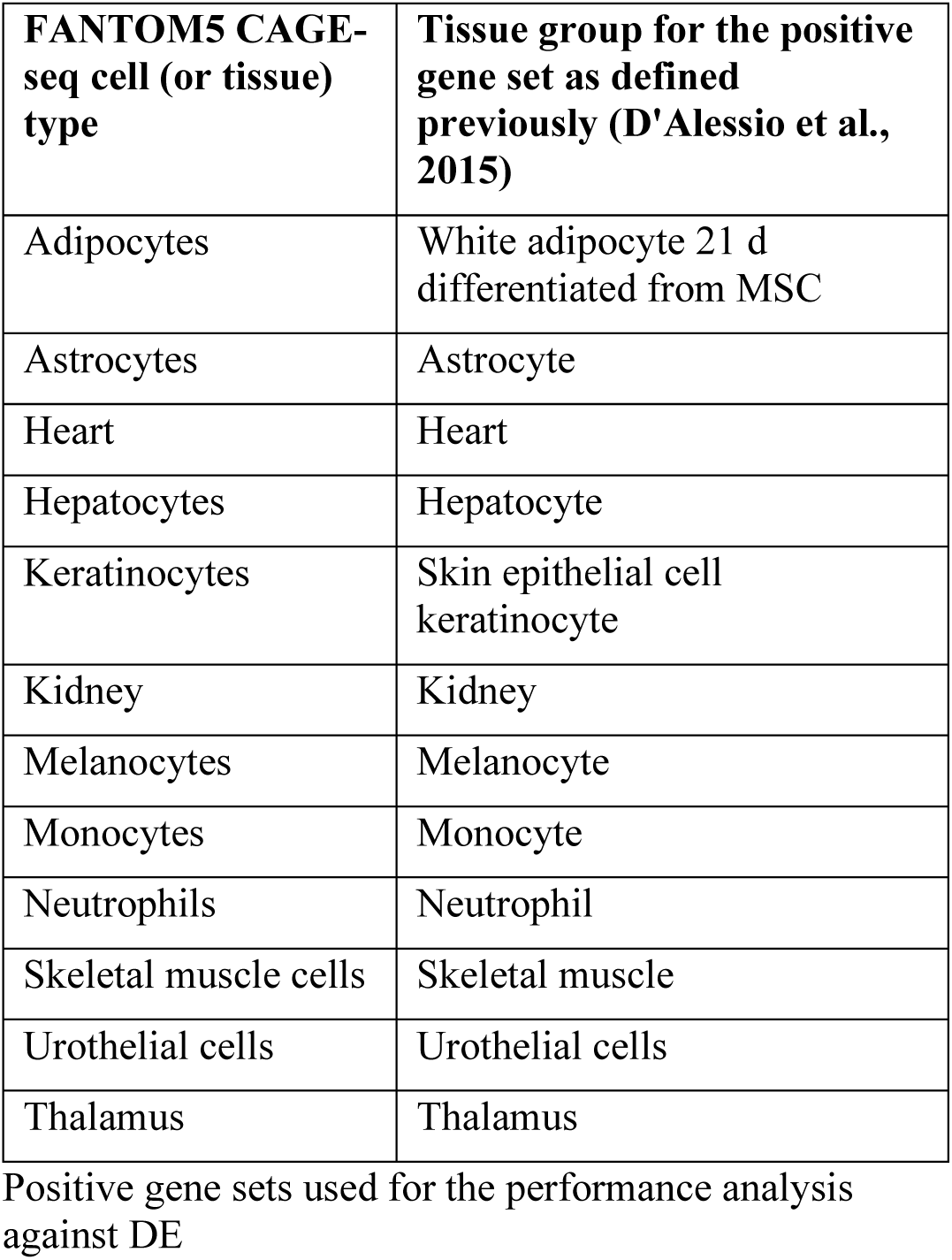

| FANTOM5 CAGE-seq cell (or tissue) type | Tissue group for the positive gene set as defined previously (D'Alessio et al., 2015) |
| --- | --- |
| Adipocytes | White adipocyte 21 d differentiated from MSC |
| Astrocytes | Astrocyte |
| Heart | Heart |
| Hepatocytes | Hepatocyte |
| Keratinocytes | Skin epithelial cell keratinocyte |
| Kidney | Kidney |
| Melanocytes | Melanocyte |
| Monocytes | Monocyte |
| Neutrophils | Neutrophil |
| Skeletal muscle cells | Skeletal muscle |
| Urothelial cells | Urothelial cells |
| Thalamus | Thalamus |

Positive gene sets used for the performance analysis against DE

### Statistical properties of the discordance score

We defined a statistical test to gauge if a DS assigned to a gene is higher than expected. The expected DS of a gene is based on two components: the expected RTS given the observed level of expression, and the expected level of expression given the observed RTS of the gene. The test first estimates *p*-values for the two test statistics, which are then combined by Fisher’s method to yield a single *p*-value based on the chi-squared distribution. In practical terms, an empirical null distribution is generated from repeated, random permutation of each test statistic of *n* comparable genes. Comparable genes are defined as those with the closest value to the gene of interest, in terms of the parameter that is *not* the test statistic (i.e. expression level or RTS). The *p*-value is then the probability that the rank of the gene of interest improves as a result of the random permutation.

### Functional enrichment analysis

To compare efficacy of TRIAGE against other existing methods in prioritizing genes functionally important to a given tissue or cell state, we used a simple systematic approach to analyze enrichment of a selected GO term. For annotation purposes, we ranked only protein-coding genes. Ranked genes by a given metric (e.g. DS, gene expression value etc.) were first binned into a percentile bin (i.e. each bin includes 1% of the total gene populations in a given dataset). At each rank bin, we quantified enrichment of a selected GO biological process (BP) term using Fisher’s exact test (one-tailed). Essentially, we tested how significantly the GO term was enriched among genes above a rank bin *m* (i.e. above top *m*% of the gene population) compared to the rest using a sliding window. The resultant significance (−*lda*_10_*p*) was plotted against the rank bin, allowing visualization of how the enrichment changes as more lowly ranked genes were included in the analysis. To identify significantly (FDR<0.05) enriched GO BP terms among top ranked genes, we used a R package topGO (Alexa, 2019).

### Consistency of the RTS between different peak callers

To assess how the RTS changes with different peak calling methods, we independently calculated the RTS using peaks identified by 3 different peak callers, namely MACS2, SPP and Homer (Feng et al., 2012; Heinz et al., 2010; Kharchenko et al., 2008). Briefly, we first downloaded mapped ChIP-seq reads in tagAlign format for the 111 Roadmap cell types. Peaks were identified by comparing H3K27me3 tag signals with the input for each peak caller across the cell types. For MACS2 peak caller, ‘callpeak’ program was used with ‘broad’ and ‘broad-cutoff’ of 0.1 options to capture broad deposition of H3K27me3 (Feng et al., 2012). For SPP peak caller, ‘get.broad.enrichment.clusters’ function (available under ‘spp’ R library) with window.sizes=1000 and z.thr=3 was used as recommended for capturing broad HMs (Kharchenko et al., 2008). For Homer peak caller, ‘findPeaks’ program was used with ‘-style histone’. This parameter ensures the peak caller to initially find peaks of size 500 bp and subsequently stitch into regions of 1000bp; a suitable approach to identify broad regions of histone modifications (Kharchenko et al., 2008). From outputs of each peak caller, we calculated RTS values as described above. Subsequently, DSs were computed for 3 distinct Roadmap tissue groups; Left ventricle (E095), Germinal matrix (E070) and T helper naïve cells (E038), using these 3 different versions of the RTS. TFs with a selected GO term (Heart development GO:0007507, Brain development GO:0007420 and T cell differentiation GO:0030217 respectively) were used as the positive gene set for the enrichment analysis.

### Accuracy of estimated RTSs

We estimated the RTS from the 111 NIH Roadmap tissue or cell types. To address potential sampling bias, we performed a bootstrapping analysis by randomly re-sampling cell types 10,000 times. For each re-sampling, we calculated the RTS for each gene. We collected the empirical bootstrap distribution of RTSs for each gene. RTSs of all 26,833 genes were estimated within 1 standard deviation of their respective mean, supporting consistency of the estimated RTS.

### Saturation of H3K27me3 signals

To understand whether the 111 cell types in Roadmap provide sufficient data to estimate stable RTSs, we developed an iterative process to quantify stability of the RTS with a differing number of cell type samples. Suppose we have *n* number of genes each of which has a RTS calculated based on H3K27me3 data observed in *k* number of cell types. We defined *saturation state* as a state where any change in the RTS for a given gene as a result of an addition of *l* number of cell types is within an arbitrarily defined range.

If the signal is in the saturation state, adding *l* number of different cell types would not result in a noticeable change to the resultant RTS. To help quantification of the RTS change, we define a term *stably ranked gene*. Suppose gene *i* has an estimated RTS derived from *k* number of cell types and is ranked at a certain position (*u*_*i,k*_). We say the gene is *stably ranked* if a resultant RTS re-calculated with an addition of *l* number of cell types put the gene at a rank position (*u*_*i,k*+*l*_) that is within a certain range of rank positions (*θ*) from the previous rank position (*u*_*i,k*_). In other words, gene *i* is stably ranked if the resultant RTS change is not large enough to shift its rank position more than *θ*. Formally, a set of *stably ranked genes* with an addition of *l* number of cell types to *k* number of cell types (*G*_*k,l*_) is a subset of all genes (*G*_*all*_) and can be written as the follow.

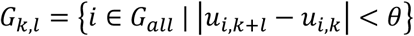

For instance, if gene A is ranked at *q* position by the RTS calculated using *k* number of cell types, the gene A is a *stably ranked gene* if the rank does not change more than *θ* positions (e.g. 1% of the total gene number) from *q* when an additional *l* number of cell types is included for the subsequent calculation.

We started with *k*=3 randomly selected cell types and iteratively calculated the proportion of *stably ranked genes* at an increment of *l*=3 randomly selected cell types without replacement until all 111 cell types were used. To address potential sampling bias, we repeated this process 1,000 times and obtained mean values and the 95% confidence interval of the estimation (**Figures S2F-G**). We used a range of different thresholds (i.e. 1∼5% of the total gene number) for the stably ranked gene (**Figure S2G**).

### Correlation between the expression and the H3K27me3 domain breadth

H3K27me3 represses transcription of the target gene but it is not known whether cell type-specific regulatory genes have a distinct functional relationship with this repressive HM. Based on strong association of cell type-specific TFs with broad H3K27me3 domains, we questioned whether the repressive effect of H3K27me3 was more prominent among the cell type-specific TFs. To this end, we first identified five classes of genes (i.e. (i) 634 VETFs to represent cell type-specific regulatory genes, (ii) 7,445 variably expressed non-TFs, (iii) 18,708 protein-coding genes, (iv) 793 non-VETFs and (v) 3,805 housekeeping genes). For each gene, we calculated a Pearsons’s correlation coefficient between the gene expression value and the breadth of the corresponding H3K27me3 domain across the 46 Roadmap cell types.

### Functional relationship between the repressive tendency and gene transcription

To understand association of RTS with the transcriptional outcome, we first ranked genes in a descending order of the RTS. We only considered coding genes and assigned them into bins of 100 genes. For the gene expression level, an average expression value of the gene set at each rank bin was calculated across the 46 cell types **(Figure 2D)**. For the expressional specificity, a proportion of cell types where a given gene was expressed (RPKM>1 or equivalent) out of the 46 cell types was calculated **(Figure 2C)**. As there were 100 genes in each rank bin, we calculated the average proportion of the 100 genes in each bin.

### Comparison of clustering accuracy

We compared clustering accuracy based on expression values and DSs using the scRNA-seq mixology benchmarking data set (Tian et al., 2019). The data set we used (*sc_10x*) was generated on the 10X platform and provided expression values as counts for each of three distinct, labelled, human lung adenocarcinoma cell lines. We assessed the clustering of this data across three methods: *SC3* (1.10.0), *CORE* (ascend 0.9.6) and *Seurat* (2.3.0) (Butler et al., 2018; Kiselev et al., 2017; Senabouth et al., 2019). Each of the clustering methods were used as detailed by their authors in their documentation or tutorials, including any filtering and scaling steps. In the SC3 clustering, the k parameter (ks) was set to 3, and the remainder of the parameters across all three algorithms were chosen as specified in their documentation or left at default.

To quantify the performance of the clustering methods, we used the Adjusted Rand Index as calculated by the *mclust* package (5.4.2) (mclust::adjustedRandIndex) (Scrucca et al., 2016), comparing the cluster assignment in each clustering method to the three cell line labels from the mixology data set. The PCA plots were generated using the ascend package (ascend::plotPCA) (Senabouth et al., 2019)

### Benchmarking TRIAGE in multi-omics platforms

We hypothesized that TRIAGE would be applicable to any quantifiable genomic data that reasonably reflects the expression level of genes. To demonstrate applicability of TRIAGE to various multi-omics platforms, we used GTEx (v8) transcriptomes (Lonsdale et al., 2013), single-cell transcriptomes (Cao et al., 2019; Friedman et al., 2018; Schaum et al., 2018), FANTOM5 cap analysis of gene expression (CAGE) peaks (Forrest et al., 2014), human proteomes (Kim et al., 2014) and Roadmap H3K36me3 ChIP-seq data (Kundaje et al., 2015). For each sample, we identified top 100 genes ranked by TRIAGE and significantly enriched GO BP terms (Benjamini-Hochberg FDR<1e-6, hypergeometric test). This was compared with the enrichment from top 100 most highly expressed genes. We extracted terms that were significantly enriched (Benjamini-Hochberg FDR<1e-6, hypergeometric test) specifically in less than a third of all comparing samples. In addition to the GO term, we provided summary plots for enrichment of the VETFs by including the lowest p-value when the significance was calculated across the percentile rank positions (**Figures 5G-H, J**).

The Tabula Muris mouse single-cell RNA-seq data encompass nearly 100,000 cells from 20 different tissue types (Schaum et al., 2018). We averaged expression values of genes for each tissue type to calculate corresponding DSs. For the CAGE data set (Forrest et al., 2014), we used the normalized CAGE tag density for the expression value of the corresponding gene. The highest CAGE tag density assigned to the gene was used so that each gene was annotated with a value from a TSS with the highest value. We selected 329 FANTOM5 samples that covers 25 distinct cell types without a disease annotation **(Table S1)**. For human proteomic data (Kim et al., 2014) covering 30 different tissue groups, we used the protein expression value to link to the corresponding gene for the quantification. For H3K36me3 data set, we collected mapped reads for H3K36me3 ChIP-seq for the 111 Roadmap cell types. For each cell type, we quantified the read density for each gene by calculating the number of reads per base-pair mapped to the RefSeq gene body. We then used the tag density as a proxy for the transcriptional abundance of the gene to calculate the DS.

To provide further validation on our analysis, we used tissue specific TFs from 233 distinct tissue groups as another positive gene sets collected from an independent study (D’Alessio et al., 2015). We extracted top 20 most tissue specific TFs (by specificity score) across the tissue groups and analyzed their enrichment among the top 100 genes ranked by TRIAGE or the expression value.

### Visualization of multi-omics datasets

To visualize performance of TRIAGE on multiple samples on a single plot simultaneously, we summarized the performance for each sample by (i) the most significant *p*-value (*y*-axis, Fisher’s exact test for enrichment of VETFs, one-sided) and (ii) the corresponding rank position where (i) is observed (*x*-axis, as the percentile rank position with 1 the highest and 100 the lowest) (**Figures 5G, 5J**). Each sample was represented as a single data point on the plot.

### Inter-species application of the TRIAGE

Given a high level of evolutionary conservation for the PRC2 (Margueron and Reinberg, 2011), we hypothesized that the RTS calculated from the human data could be effectively applied to equivalent genes of other species. To test this, we first downloaded a range of transcriptomic data sets from different species covering the 5 selected tissue-groups (**Table S1**). We then performed inter-species gene mapping using online Ensembl bioMart (http://asia.ensembl.org/biomart/martview/) by identifying human orthologues (Haider et al., 2009). Only genes mappable to human orthologues were included in the analysis.

### Biological effect of perturbing TRIAGE priority genes in hESCs

To evaluate the biological significance of perturbing TRIAGE priority genes, we utilized public resource data (Nakatake et al., 2020) in which 714 doxycycline-inducible transgene (including 418 TFs) overexpression (OE) hESC lines were established and assessed for transcriptome changes via RNA-seq/microarray 48hrs after the presence or absence of dox (Figure S2H). A total of 510 transgene OE cell lines were sequenced, out of which we identified 145 TRIAGE priority genes and 363 non-priority genes, excluding DUX4 and LHFPL6 which did not have RTS scores (Figure S2E). Within these two groups, the number of differentially expressed genes after induction of a transgene (comparing control vs. +dox samples for each cell line) was determined using ExAtlas (>2-fold-change, Benjamini-Hochberg FDR<0.05) (Sharov et al., 2015). The total number of DEGs was compared between TRIAGE priority (n=145) and non-priority (n=363) groups using an unpaired t-test with Welch’s correction (*p*<0.0001, two-tailed).

### Melanoma gene set enrichment

We used published pre-processed single cell RNA-seq data from melanoma tumors (Tirosh et al., 2016).1,252 melanoma cells were isolated from the set of approximately 4,000 cells based on the authors annotations. Melanoma proliferative and invasive gene sets were obtained from the same source. Based on the ratio of the average gene expression of the proliferative to invasive genes; the top 50 most proliferative and the top 50 most invasive cells were identified. TRIAGE was applied to the melanoma gene expression profiles to produce DS profiles. Averages of both expression and the discordance profiles of the top 50 proliferative and invasive cells were taken. Genes were subsequently ranked by the expression and discordance values for these representative profiles. For the ranked genes, fishers exact test was iteratively performed from the top ranked genes down the list as described above; adding 1% of the genes at each iteration. Gene sets tested for enrichment were obtained from published melanoma data sets (Tirosh et al., 2016). False discovery rate was used to correct for multiple hypothesis testing.

### Heart Failure pathogenesis dataset

To determine the utility of TRIAGE in identifying regulatory elements and processes of disease in heart failure pathogenesis we used published pre-processed bulk RNA-seq data from adult mouse ventricles (GSE58453 and GSE68509) (Duan et al., 2017). Briefly, heart failure was induced using transverse aortic constriction (TAC) and the small molecule pan-BET inhibitor JQ1 was used to treat the TAC condition. TRIAGE was applied to the gene set of each condition (SHAM, TAC, TAC+JQ1) to produce DS profiles. Genes were subsequently ranked by the expression and discordance values for these representative profiles. For the ranked genes, Fishers exact test was iteratively performed from the top ranked genes down the list for GO terms of interest.

| Candidate genes | RTS | Population of interest | Dataset | Efficient CRISPR K/D or K/O |
| --- | --- | --- | --- | --- |
| SIX3 | 0.539 | Definitive Endoderm/Mesendoderm | hiPSC cardiac differentiation, Day 2 | Yes |
| GAD1 | 0.176 | Endothelial Cells | hiPSC cardiac differentiation, Day 5 | Yes |
| CRLF1* | 0.079 | Mesendoderm | hiPSC cardiac differentiation, Day 5 | No |
| ZNF503* | 0.077 | Cardiac Progenitor Cells | hiPSC cardiac differentiation, Day 5 | No |
| RNF220 | 0.304 | First Heart Precursors | Ciona heart development | Yes |
| IRX4/6 | 0.318/<br>0.179 | Atrial Siphon Muscle | Ciona heart development | Yes |
| TBX2/3 | 0.188/<br>0.332 | Trunk Ventral Cells | Ciona heart development | No |
| IGF1/2 | 0.001/<br>0.164 | First Heart Precursors | Ciona heart development | No |
| IGF2BP1/2/3* | 0.172 | First Heart Precursors | Ciona heart development | No |

### Gene Ontology Visualization

Gene ontology analysis was performed using DAVID with significance threshold set at FDR<0.05. The *p*-values from gene ontology analysis were visualized with the radius of the circle proportional to the negative natural log of the input *p*-value.

### iTranscriptome Sample Preparation and Data Analysis

Samples were generated according to the methodology previously published (Peng et al., 2016). E5.5, E6.0, E6.5, E7.0 and E7.5 embryos were cryo-sectioned along the proximal-distal axis. Populations of approximately 20 cells were collected from different regions of the cross-section by laser microdissection and processed for RNA sequencing (see **Figure S9E**). From the RNA-sequencing dataset, differentially expressed genes (DEGs) were screened first by unsupervised hierarchical clustering method to group samples in the respective regions. Genes with an expression level FPKM>1 and a variance in transcript level across all samples greater than 0.05 were selected. To identify inter-region specific DEGs, each of these selected genes was submitted to a t-test against the level of expression in the other regions. Genes with a *p*-value<0.01 and a fold change>2.0 or <0.5 were defined as DEGs. The gene expression pattern (region and level of expression by transcript reads) of the gene of interest was mapped on the corn plots, where each kernel represents the cell population sampled at a defined position in the tissue layers of the embryo, to generate a digital 2D rendition of the expression domain that emulated the display of the result of whole mount *in situ* hybridization.

### Selection of candidate genes for biological validation using TRIAGE

Candidate genes for biological validation of TRIAGE predictions in *in vitro* hiPSC and *in vivo* Ciona model systems were selected using the following criteria. Genes were selected if (1) ranked in the top 10 TRIAGE-expression gene list (2) fall within the top 1359 TRIAGE priority genes (3) have no known role in the population of interest (4) transcription factor or signaling molecule, and (5) efficient CRISPRi knockdown in hPSCs or CRISPR knockout in Ciona (Figure S9A). Using this selection criteria the following genes were selected for genetic loss of function:

*Note: CRLF1, ZNF503 and IGF2BP1/2/3 were ranked within the top 25 TRIAGE-expression list, and were chosen based on high population specificity, whilst all other genes fell within the top 10.

For these genes CRISPR K/D hPSC cell lines and CRISPR K/O Ciona embryos were generated testing 3 gRNAs per gene. Efficiency of genetic loss of function was then tested via qPCR in hPSCs and peakshift assay in Ciona. Of the genes which displayed efficient loss of function, SIX3 and RNF220 were chosen for further biological phenotype validation because they had higher RTSs and had no known roles in the relevant populations of interest.

### Generation and Maintenance of Human ESC/iPSC Lines

All human pluripotent stem cell studies were carried out in accordance with consent from the University of Queensland’s Institutional Human Research Ethics approval (HREC#: 2015001434). WTC CRISPRi GcaMP hiPSCs (Karyotype: 46, XY; RRID CVCL_VM38) were generated using a previously described protocol(Mandegar et al., 2016) and were generously provided by M. Mandegar and B. Conklin (UCSF, Gladstone Institute). WTC CRISPRi SIX3-g2 hiPSCs were generated in this study (see below). All cells were maintained as previously described(Palpant et al., 2017a). Briefly, cells were maintained in mTeSR media with supplement (Stem Cell Technologies, Cat.#05850) at 37° C with 5% CO_2_. WTC CRISPRi GCaMP and WTC CRISPRi SIX3-g2 hiPSC lines were maintained on Vitronectin XF (Stem Cell Technologies, Cat.#07180) coated plates.

### WTC CRISPRi SIX3-g2 hiPSCs

3 separate guide RNAs (gRNA) targeting the CAGE-defined transcriptional start sites of the human SIX3 sequence were designed and cloned into the pQM-u6g-CNKB doxycycline-inducible construct and transfected into WTC CRISPRi GCaMP hiPSCs using the Neon transfection system (Invitrogen, Cat.#MPK1096). For electroporation, 0.5µg DNA and 1×10^5^ dissociated hiPSCs were mixed in 10µL resuspension buffer R (Invitrogen, Cat.#MPK1096). Electroporation parameters were as follows: pulse voltage, 1300V; pulse width, 30ms; and pulse number, 1. Cells were then plated in Vitronectin XF (Stem Cell Technologies, Cat.#07180) coated plates in mTeSR media (Stem Cell Technologies, Cat.#05850) supplemented with 10µM of Y-27632 (Stem Cell Technologies, Cat.#72308). Stable clones were selected using successive rounds of re-plating with blasticidine at 10µg/ml (Sigma, Cat.#15205). Populations were tested for knockdown efficiency by qPCR following doxycycline addition at 1 µg/ml (Sigma, Cat.#D9891) continuously from day 0 of cardiac-directed differentiation (*n*=12-16 technical replicates per condition from 4-5 experiments). WTC CRISPRi SIX3-g2 line displayed high knockdown efficiency and therefore was chosen.

#### Guide RNAs designed

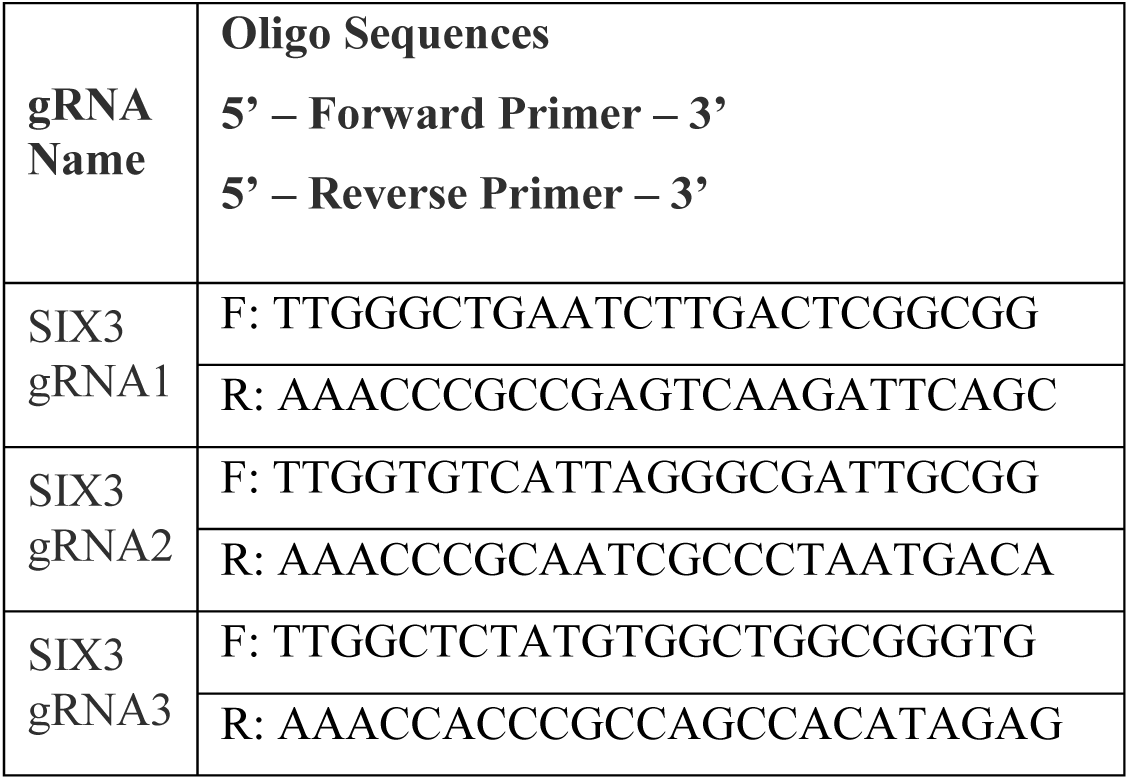

### Cell Culture

All human pluripotent stem cell studies were carried out in accordance with consent from the University of Queensland’s Institutional Human Research Ethics approval (HREC#: 2015001434). hiPSCs were maintained in mTeSR media (Stem Cell Technologies, Cat.#05850). Unless otherwise specified, cardiomyocyte directed differentiation using a monolayer platform was performed with a modified protocol based on previous reports(Burridge et al., 2014). On day −1 of differentiation, hPSCs were dissociated using 0.5% EDTA, plated into vitronectin coated plates at a density of 1.8 x 10^5^ cells/cm^2^, and cultured overnight in mTeSR media. Differentiation was induced on day 0 by first washing with PBS, then changing the culture media to RPMI (ThermoFisher, Cat.#11875119) containing 3µM CHIR99021 (Stem Cell Technologies, Cat.#72054), 500µg/mL BSA (Sigma Aldrich, Cat.#A9418), and 213µg/mL ascorbic acid (Sigma Aldrich, Cat.#A8960). After 3 days of culture, the media was replaced with RPMI containing 500µg/mL BSA, 213µg/mL ascorbic acid, and 5µM Xav-939 (Stem Cell Technologies, Cat.#72674). On day 5, the media was exchanged for RPMI containing 500µg/mL BSA, and 213µg/mL ascorbic acid without supplemental cytokines. From day 7 onwards, the cultures were fed every 2 days with RPMI plus 1x B27 supplement plus insulin (Life Technologies Australia, Cat.#17504001).

### Quantitative RT-PCR

For quantitative RT-PCR, total RNA was isolated using the RNeasy Mini kit (Qiagen, Cat.#74106). First-strand cDNA synthesis was generated using the Superscript III First Strand Synthesis System (ThermoFisher, Cat.#18080051). Quantitative RT-PCR was performed using SYBR Green PCR Master Mix (ThermoFisher, Cat.#4312704) on a ViiA 7 Real-Time PCR System (Applied Biosystems). The copy number for each transcript is expressed relative to that of housekeeping gene *HPRT1*. Samples were run in biological triplicate. FC was calculated on a gene by gene basis as gene expression divided by control gene expression. The following are qRT-PCR primers utilized in this study:

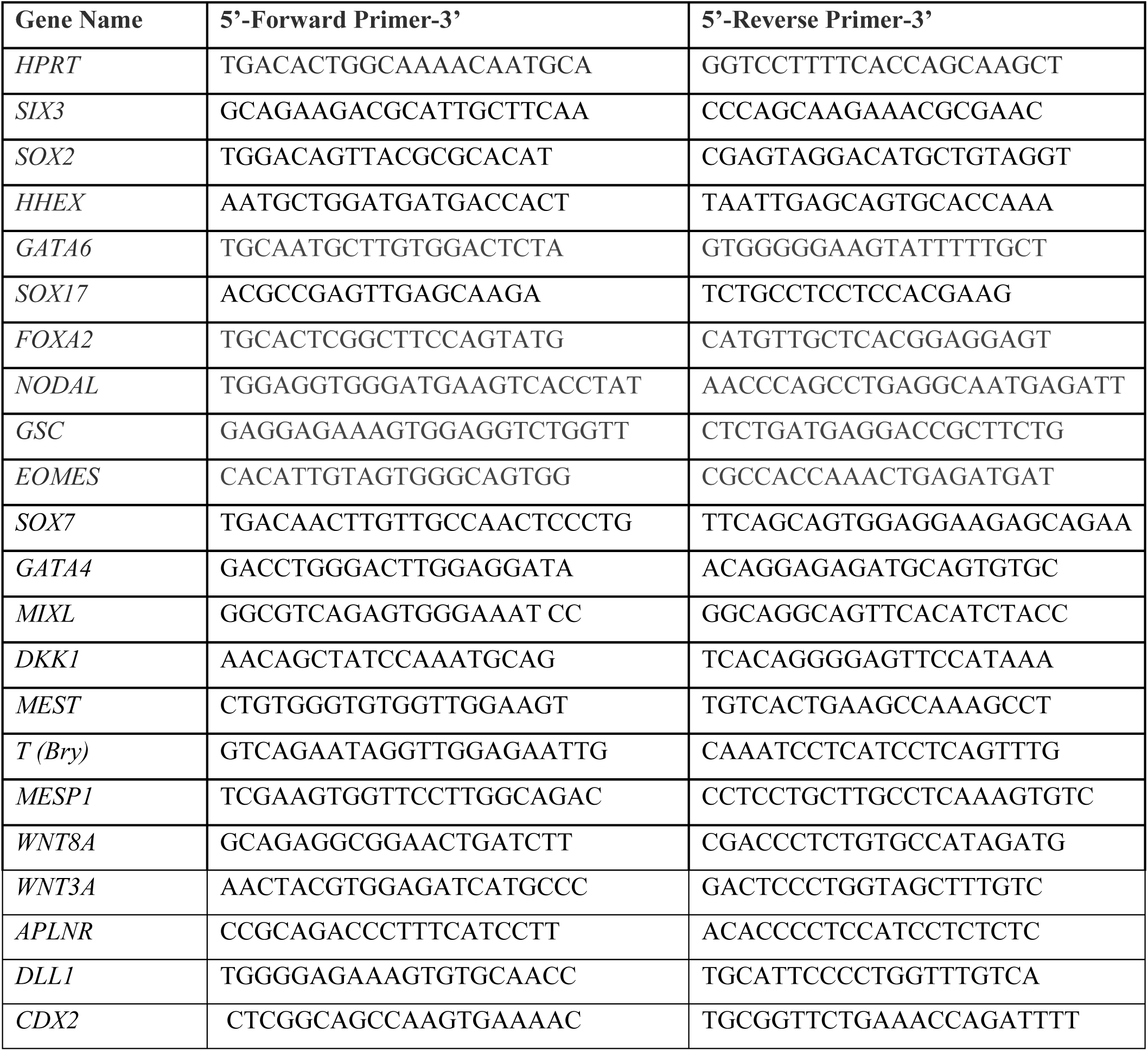

### Flow Cytometry

On day 2 of cardiac differentiation cells were dissociated using 0.5% EDTA and put into blocking buffer of 50% fetal bovine serum (FBS) in Dulbecco’s Modified Eagle Medium (DMEM)/F12 (ThermoFisher, Cat.#11320033). Cells were then pelleted and resuspended in 10% FBS in DMEM media. Cells were labeled live for flow cytometry using CD184 (BD Biosciences, Cat.#555974), EpCAM (BD Biosciences, Cat.#347199) and corresponding isotype controls were used to gate the cells. On day 15 of cardiac differentiation cells were fixed with 4% paraformaldehyde (Sigma, Cat.#158127) and permeabilized in 0.75% saponin (Sigma, Cat.#S7900). On day 15 of cardiac differentiation fixed cells were labeled for flow cytometry using alpha-actinin (Miltenyi Biotec, Cat.#130106937) and corresponding isotype control. Cells were analyzed using a BD FACSCANTO II (BD Biosciences) with FACSDiva software (BD Biosciences). Data analysis was performed using FlowJo (Tree Star).

### *Ciona robusta* CRISPR/Cas9 gene editing

For Rnf220 (KH2012:KH.C8.791) loss of function, 3 sgRNAs were designed to avoid genomic off-targets and tested as described (Stolfi et al., 2014). sgRNA expressing cassettes (U6 > sgRNA) were assembled by single step overlap PCR. Individual PCR products (∼25 µg) were electroporated with EF1a > NLS::Cas9::NLS (20 µg), Myod905 > Venus (50 µg), driving ubiquitous expression of Cas9 and a widely expressed fluorescent reporter construct, respectively, as described (Christiaen et al., 2009). Efficient electroporation was confirmed by observation of fluorescence before genomic DNA extraction around 16 hpf (18°C) using QIAamp DNA Micro kit (Qiagen, German Town, MD). Mutagenesis efficacy of individual sgRNAs, as a linear function of Cas9-induced indel frequency, was estimated from electrophoregrams following Sanger sequencing of the targeted regions amplified from extracted genomic DNA by PCR. Results of the relative quantification of the indel frequency (‘corrected peakshift’ of 22% and 16%) for sgRNAs 2 and 3 was considered high enough for both sgRNAs targeting Rnf220, which were finally selected. The corresponding cassettes were cloned into plasmid for repeated electroporations to study the loss of function of Rnf220. In order to control the specificity of the CRISPR/Cas9 system, sgRNAs targeting *Tyrosinase*, a gene not expressed in the cardiopharyngeal lineage, was electroporated in parallel. For imaging experiments, sgRNAs (25 µg) were electroporated with Mesp > NLS::Cas9::NLS (20 µg), Mesp > H2B:GFP (50 µg) and Mesp > mCherry (50 µg). Sequences of the DNA targets and oligonucleotides used for the sgRNAs:

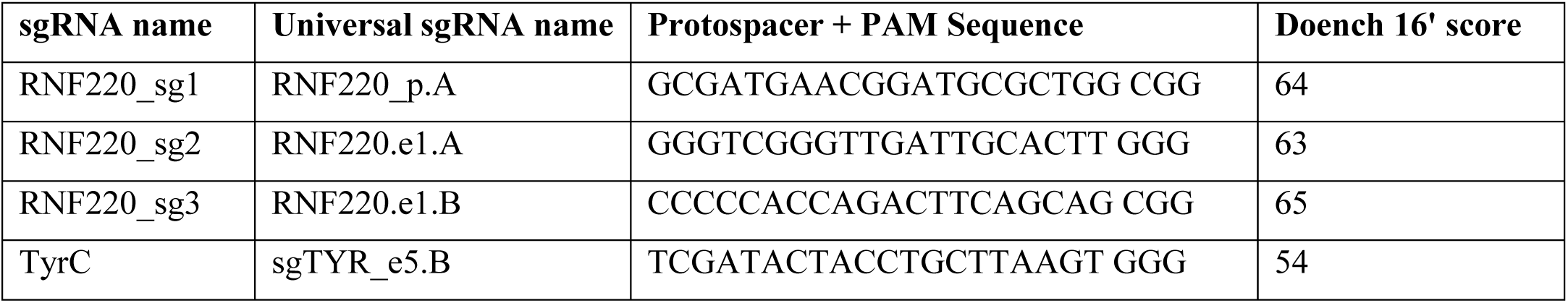

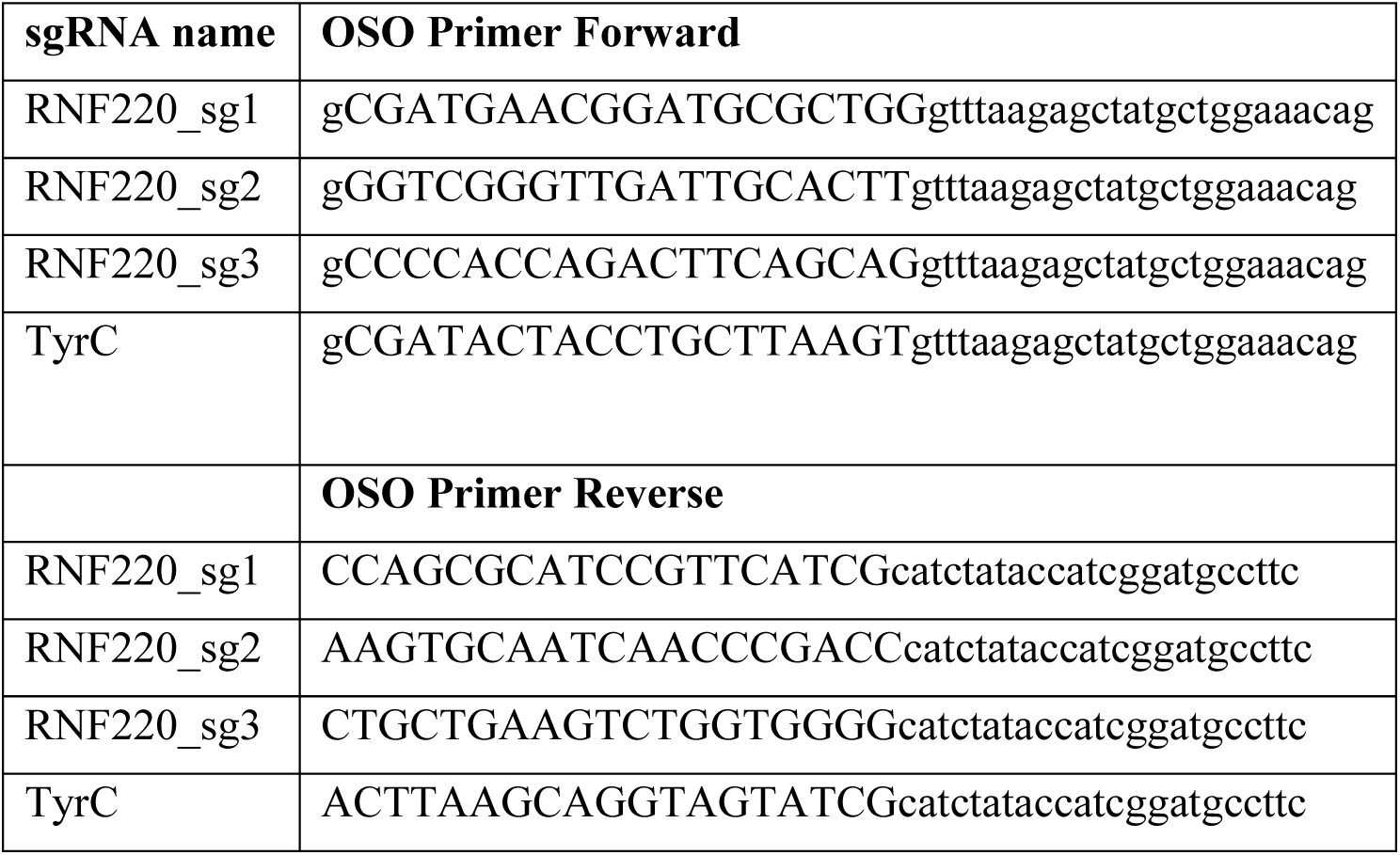

Embryos were fixed in 4% MEM-FA for 30 minutes, incubated with an NH_4_Cl solution, and imaged using Leica SP8 X Confocal microscope.

## RESOURCE AVAILABILITY

### Lead contact

Further information and requests for resources and reagents should be directed to and will be fulfilled by the Lead Contact, Nathan J. Palpant.

### Materials availability

This study did not generate new materials.

### Data and Code availability

- Source data statement: This paper analyzes existing, publicly available data. These datasets’ accession numbers are provided in the manuscript text and Supplementary Table 1.
- Code statement: We provide the TRIAGE source code written in Python and R (https://github.com/woojunshim/TRIAGE). Users can also run the TRIAGE analysis using a web accessible interface (http://bioinf.scmb.uq.edu.au/adhoc/).
- Scripts statement: The scripts used to generate the figures reported in this paper are available at https://github.com/woojunshim/TRIAGE.

Any additional information required to reproduce this work is available from the Lead Contact.

## QUANTIFICATION AND STATISTICAL ANALYSIS

Unless otherwise noted, all data are represented as mean ± standard error of mean (SEM). Indicated sample sizes (*n*) represent biological replicates including independent cell culture replicates and individual tissue samples. No methods were used to determine whether data met assumptions of the statistical approach or not. Due to the nature of the experiments, randomization was not performed and the investigators were not blinded. Statistical significance was determined in GraphPad Prism 7 software by using student’s t test (unpaired, two-tailed) or ordinary one-way ANOVA unless otherwise noted. Results were considered to be significant at *p* < 0.01(*). Statistical parameters are reported in the respective figures and figure legends. All statistical data are represented as mean ± SEM.

## ACKNOWLEDGEMENTS

E.S acknowledges funding by Children’s Hospital Foundation Queensland (Award Reference Number: 50268). B.V. acknowledges funding by American Heart Association grant #18PRE33990254. The *Ciona* work was supported by NIH/NHLBI award R01 HL108643 to L.C. M.A. was supported by the Swiss National Science Foundation (project P2LAP3_178056), P.P.L.T. is supported by the National Health and Medical Research Council of Australia (Grant 1110751). N.P is supported by the National Health and Medical Research Council of Australia (Grant 1143163) and the Australian Research Council (Grant SR1101002).

## AUTHOR CONTRIBUTIONS

**WJS**: Developed the computational basis for the study, performed data analysis and wrote the manuscript

**ES**: Contributed to experimental and computational design for the study, performed data analysis, carried out functional genetic studies in hPSCs and wrote the manuscript

**JX**: Assisted with computational analysis and developed web interactive interface

**MA**: Performed computational analysis on HF pathogenesis data

**GA**: Performed computational analysis on HF pathogenesis data

**SS:** Assisted the computational analysis on different single-cell data platforms

**BB**: Performed computational analysis on melanoma studies

**YS**: Performed computational analysis on Mouse Organogenesis Cell Atlas data

**CB:** Contributed EpiMap data

**BV**: Performed functional analysis on *ciona* and validated the findings

**GP**: Assisted with spatiotemporal transcriptomic profiling of mouse gastrulation

**NJ**: Assisted with spatiotemporal transcriptomic profiling of mouse gastrulation

**YW:** Helped with computational analysis of epigenetic data

**MK:** Contributed EpiMap data

**MP**: Assisted with analysis and interpretation of melanoma data

**AS:** Carried out experiments involving melanoma analysis

**PT**: Supervised work on spatiotemporal transcriptomic profiling of mouse gastrulation

**LC**: Performed functional analysis on *ciona* and validated the findings

**QN:** Provided assistance to implement TRIAGE on single-cell data sets

**MB** and **NJP**: Supervised the project, raised funding, and wrote the manuscript

