## Supplementary Material for "Conserved epigenetic regulatory logic infers genes governing cell identity"

### SUPPLEMENTAL FIGURES

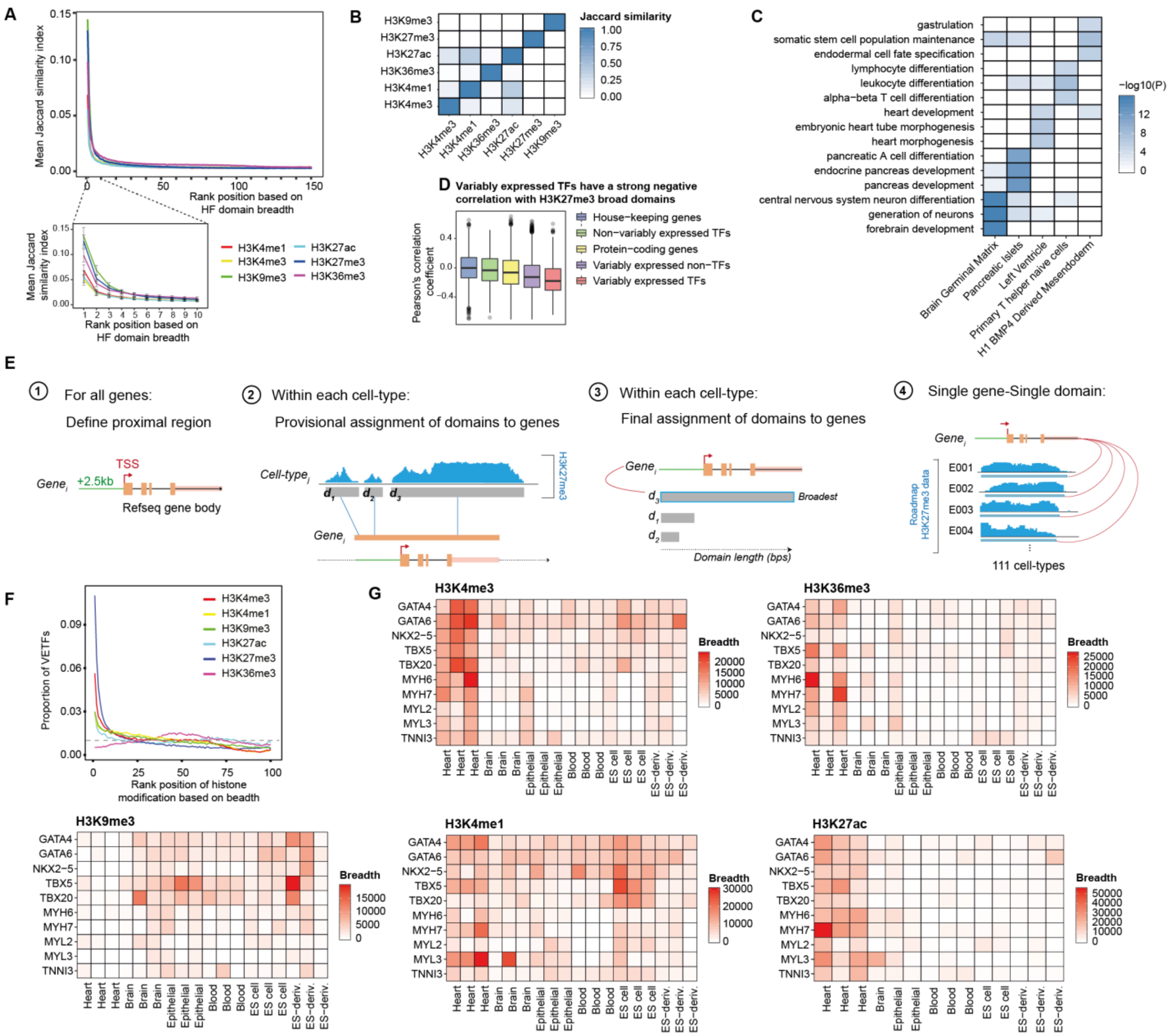


**Figure S1. H3K27me3 histone modification (HM) domains have distinct functional association with cell type-specific regulatory genes (Related to Figure 1).**

**(A)** Similar genes are associated with broad HM domains across the Roadmap tissue and cell types. Genes are ordered by the breadth of the associated HM domain and grouped into bins of 100 genes (x-axis) and average Jaccard similarity index of the gene bins for all possible pairs between cell types is calculated (y-axis). (Inset) Top 100 genes (i.e. first rank bin) are significantly more similar than other genes with narrower domains (*p*<2.2e-16 for all HMs, Wilcoxon rank-sum test, one-tailed). Scale bars show the 95% confidence interval.

**(B)** Jaccard similarity between top 200 genes that are most frequently associated with the broad HM. Distinct gene sets are identified by different HM types.

**(C)** Variably expressed transcription factors are cell-type specific. Enriched tissue-specific gene ontology (GO) biological process (BP) terms associated with most highly expressed 50 VETFs in 5 different tissue or cell types (Fisher's exact test, one-tailed); Brain germinal matrix (E070), Pancreatic islets (E087), Left ventricle (E095), Primary T helper naïve cells (E038) and H1 BMP4-derived mesendoderm (E004).

**(D)** Correlation between the H3K27me3 domain breadth and the corresponding gene expression value observed in 46 Roadmap tissue and cell types. Stronger negative correlation is observed for variably expressed TFs (n=634, median Pearson's *r* = -0.181), compared to variably expressed non-TFs (n=7,406, median Pearson's *r* = -0.128, *p*=2.57e-07), all protein-coding genes (n=18,490, median Pearson's *r* = -0.064, *p*=2.55e-25), non-variably expressed TFs (n=793, median Pearson's *r* = -0.035, *p*=2.31e-23) and housekeeping genes (n=3,818, median Pearson's *r* = -0.002, *p*=7.7e-54) (Welch's t-test, one-tailed).

**(E)** Schematic of the method for histone modification domain assignment to genes in 111 NIH Epigenome Roadmap samples. See Methods for detailed description of steps.

**(F)** VETFs are enriched in H3K27me3 broad domains. Distribution of VETFs in gene bins ranked by the HM breadth. Each bin contains 1% of 26,833 RefSeq genes. At top 1% rank position, H3K27me3 are more frequently associated with VETFs across Roadmap cell types (median VETF count=72), compared to the other HMs; H3K4me1 (17 VETFs, *p*=3.36e-35, Wilcoxon-rank sum test), H3K4me3 (33 VETFs, *p*=2.22e-22), H3K9me3 (19 VETFs, *p*=2.72e-35), H3K27ac (18 VETFs, *p*=6.52e-31) and H3K36me3 (3 VETFs, *p*=4.96e-38). Average proportion of VETFs in a given rank bin is shown in the plot. Dashed line shows the uniform distribution (proportion=0.01).

**(G)** Cardiac transcription factors and structural genes are not distinguished by any histone modification aside from H3K27me3 broad domains. Breadths of HM domains associated with selected cardiac-specific regulatory (*GATA4, GATA6, NKX2-5, TBX5, TBX20*) and structural (*MYH6, MYH7, MYL2, MYL3, TNNI3*) genes in 18 Roadmap samples; Heart (E095, E104, E105), Brain (E070, E071, E082), Epithelial (E057, E058, E059), Blood (E037, E038, E047), ES cell (E003, E016, E024) and ES-deriv. (E004, E005, E006).

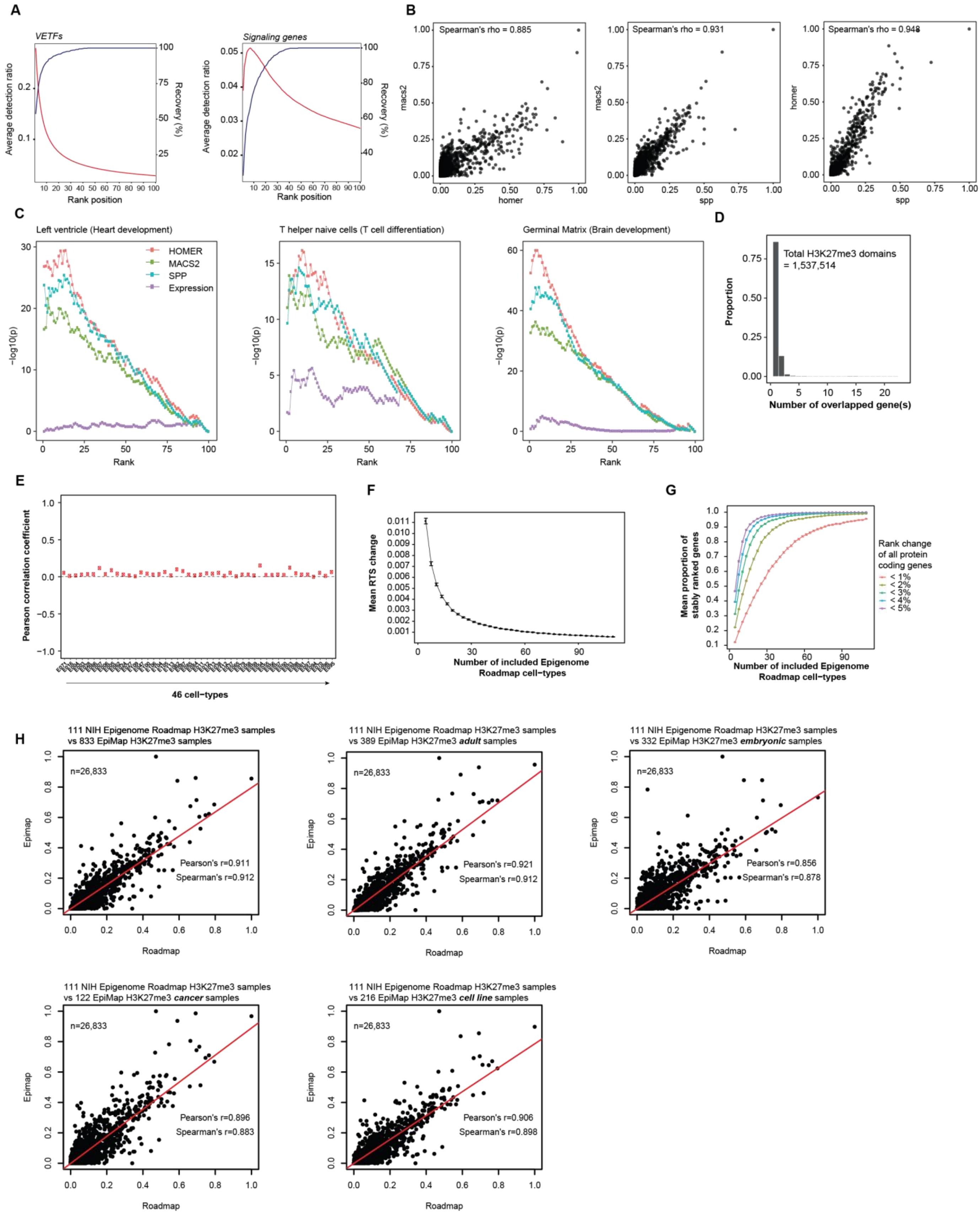


**Figure S2. Repressive tendency scores (RTS) are stable and reproducible (Related to Figures 2 and 3).**
**(A)** VETFs and signaling genes are strongly enriched in the broadest domains of H3K27me3. Data show sensitivity of the H3K27me3 breadth to recover VETFs and KEGG signaling genes. Genes are sorted by the breadth of the assigned H3K27me3 domain and binned into a rank position (equivalent to 1% of all genes, n=26,833). At each rank position in each Roadmap cell type, detection ratio (i.e. positive hits divided by the number of domains drawn) and recovery percentage (i.e. a percentage of the positive genes recovered) are calculated.

**(B)** RTS scores are highly consistent regardless of peak calling algorithm. Spearman's correlation coefficients between RTS variants based on H3K27me3 identified by 3 different peak callers (i.e. MACS2, SPP2 and HOMER), with parameters optimized for capturing broad depositions of H3K27me3.
**(C)** Enrichment of cell type-specific regulatory genes by applying TRIAGE with the different RTS calculated by different peak callers. Positive genes used are TFs with (i) 'Heart development' GO:0007507 for left ventricle (E095), (ii) 'T cell differentiation' GO:0030217 for T helper naïve cell (E038), and (iii) 'Brain development' GO:0007420 for brain germinal matrix (E070).
**(D)** The number of genes overlapped with assigned H3K27me3 domains (n=1,537,514). Approximately 85% of the assigned domains overlap a single gene.
**(E)** Discordance scores are not correlated with original expression data measured across 46 Epigenome Roadmap samples. Data show Pearson's correlation coefficients between expression values and discordance scores for all protein coding genes across the 46 Roadmap cell types.
**(F)** RTS changes from a cumulative addition of Roadmap epigenomic samples by bootstrapping. The process is repeated 1,000 times. Error bars show the 95% confidence interval.
**(G)** Regardless of gene input threshold, Epigenome Roadmap samples are sufficient to calculate a stable RTS. Proportion of stably ranked genes from a cumulative addition of samples using different input gene thresholds. The process is repeated 1,000 times. Error bars show the 95% confidence interval.
**(H)** Correlation analysis of RTS values calculated from 111 NIH Epigenome Roadmap samples compared against diverse cell and tissue samples collated in the EpiMap demonstrate highly similar results for RTS calculations independent of sample input.

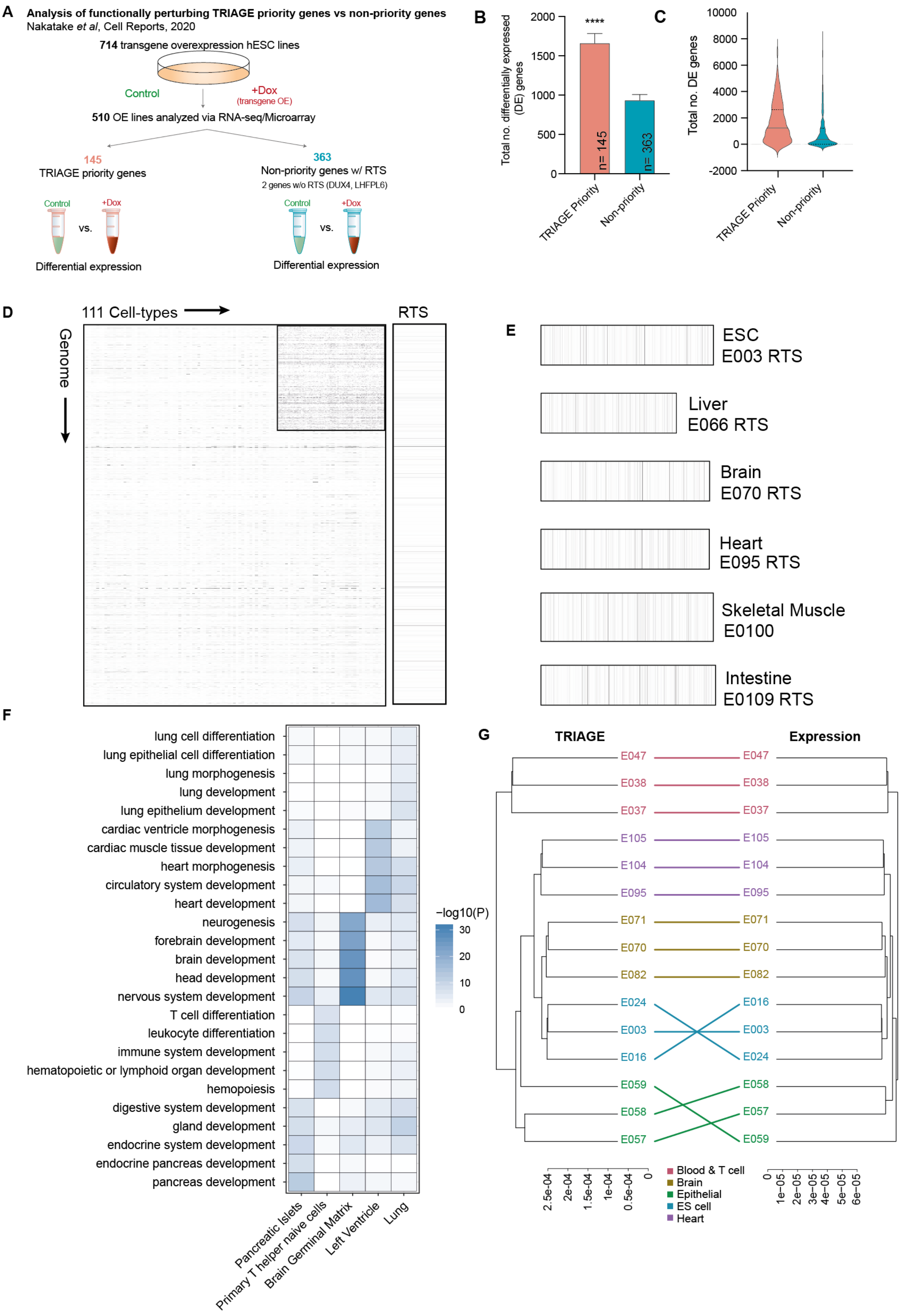
A
Analysis of functionally perturbing TRIAGE priority genes vs non-priority genes
Nakatake et al, Cell Reports, 2020
714 transgene overexpression hESC lines
Control
+Dox
(transgene OE)
510 OE lines analyzed via RNA-seq/Microarray
145
TRIAGE priority genes
363
Non-priority genes w/ RTS
2 genes w/o RTS (DUX4, LHFPL6)
Control
+Dox
vs.
Differential expression
B
Total no. differentially expressed (DE) genes
****
n= 145
n= 363
TRIAGE Priority
Non-priority
C
Total no. DE genes
TRIAGE Priority
Non-priority
D
111 Cell-types
Genome
RTS
E
ESC
E003 RTS
Liver
E066 RTS
Brain
E070 RTS
Heart
E095 RTS
Skeletal Muscle
E0100
Intestine
E0109 RTS
F
lung cell differentiation
lung epithelial cell differentiation
lung morphogenesis
lung development
lung epithelium development
cardiac ventricle morphogenesis
cardiac muscle tissue development
heart morphogenesis
circulatory system development
heart development
neurogenesis
forebrain development
brain development
head development
nervous system development
T cell differentiation
leukocyte differentiation
immune system development
hematopoietic or lymphoid organ development
hemopoiesis
digestive system development
gland development
endocrine system development
endocrine pancreas development
pancreas development
−log10(P)
Pancreatic Islets
Primary T helper naive cells
Brain Germinal Matrix
Left Ventricle
Lung
G
TRIAGE
Expression
E047
E038
E037
E105
E104
E095
E071
E070
E082
E024
E003
E016
E059
E058
E057
Blood & T cell
Brain
Epithelial
ES cell
Heart

**Figure S3. RTS priority genes identify potent regulators of cell differentiation (Related to Figures 2 and 3).**
**(A)** Schematic overview of Nakatake et al. data (Nakatake et al., 2020). To evaluate the biological significance of perturbing TRIAGE-prioritized genes RNA-seq/microarray data from 510 doxycycline-inducible transgene overexpression hESC lines were analyzed, comparing samples 48hrs after the presence or absence of dox.
**(B-C)** Total number of differentially expressed genes **(B)** and violin plot **(C)** showing overall distribution comparing control vs. +dox samples for each cell line after induction of TRIAGE priority transgenes (n=145) versus non-priority transgenes (n=363) ($p$=<0.0001, Welch's t-test, two-tailed).
**(D)** **(Left)** A heatmap showing H3K27me3 domain breadth assigned to 26,833 RefSeq genes (y-axis) across the 111 Roadmap tissue and cell types (x-axis). The band darkness shows the breadth of the assigned domain. **(Inset)** A binary heatmap showing top 5% broadest domains assigned to the genes (y-axis) across the Roadmap samples (x-axis). **(Right)** The resultant RTS values for corresponding genes (y-axis). The band darkness shows the RTS.
**(E)** Customized RTS scores based on orthologous input gene expression data from diverse cell and tissue types. TRIAGE utilizes a set of RTS values specific for the input transcriptome. The band darkness shows the RTS for expressed genes (RPKM>0) sorted by the cell-specific input expression data ordered from high expression to low expression, left to right.
**(F)** Top ranked TRIAGE genes from RNA-seq data of different tissues show cell-type specific enrichment in gene ontologies associated with organ development and morphogenesis. Data show enrichment of tissue or cell type-specific GO BP terms among top 1% genes ranked by TRIAGE in 5 distinct Roadmap tissue or cell types (Fisher's exact test, one-tailed); Pancreatic Islets (E087), Primary T helper naïve cells from peripheral blood (E038), Brain Germinal Matrix (E070), Left Ventricle (E095), Lung (E096).
**(G)** Similarity between selected 15 Roadmap samples based on (i) the original transcriptome (Expression) or (ii) the discordance score (TRIAGE). Distance between samples is 1-Pearson's correlation coefficient.

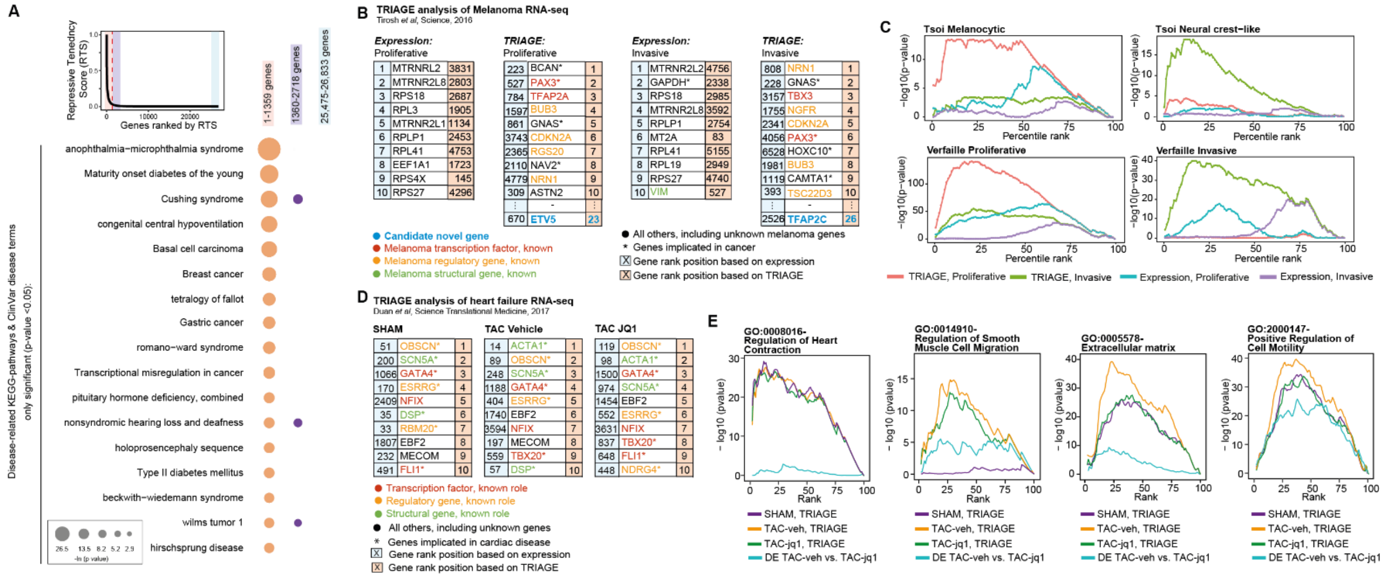


**Figure S4. TRIAGE analysis reveals enrichment of disease related gene programs (Related to Figures 2 and 3)**
**(A)** Genes with high RTS scores are enriched in disease processes. **(Top)** Distribution of the RTS values. Red dashed line is the inflection point on the interpolated curve (RTS=0.03) above which genes exhibit a substantially higher RTS than the rest ($n$=1,359, the priority genes). **(Bottom)** Functional enrichment of disease-related KEGG pathways and ClinVar disease terms in genes ranked by the RTS (Fisher's exact test, one-sided).
**(B)** Tables showing the top ranked genes from proliferative melanoma cells or invasive melanoma cells indicating rank position by original expression (left) or TRIAGE (right). Genes are identified based on their known roles as structural or regulatory genes in melanoma.
**(C)** Enrichment of positive gene sets for proliferative and invasive melanoma demonstrating high specificity of enrichment for cell type-specific gene signatures only with TRIAGE (Fisher's exact test, one-tailed).
**(D)** Top 10 genes ranked by expression (left) or TRIAGE (right) from SHAM, TAC-vehicle and TAC-JQ1 data from bulk RNA-seq of mice subjected to sham surgery (SHAM), transverse aortic constriction (TAC-vehicle) and TAC treated with JQ1 (TAC-JQ1).
**(E)** Enrichment of GO terms associated with cardiac biology and heart failure stress response mechanisms comparing each condition analyzed by TRIAGE or DE analysis (TAC-veh vs TACJQ1) (Fisher's exact test, one-tailed). Genes are ranked by either the expression value or TRIAGE and binned into a percentile bin.

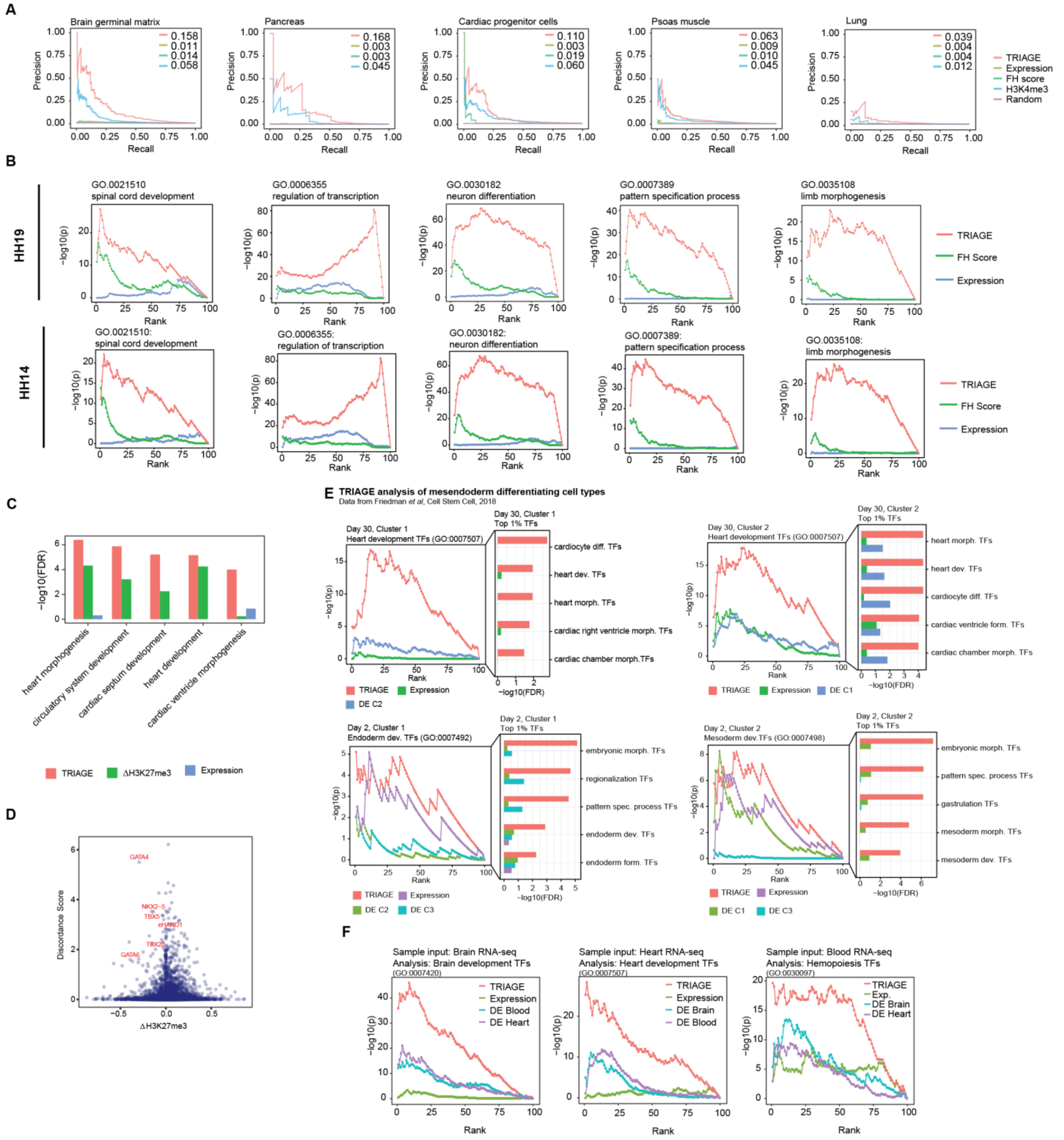


**Figure S5. TRIAGE prioritizes key regulators of cell identity across diverse cell and tissue types (Related to Figures 3 and 4).**

**(A)** TRIAGE consistently identifies regulatory genes controlling development and differentiation of diverse cell and tissue types. Data show precision-recall curves (PRC) for the 5 different tissue groups. Area under curve (AUC) values are shown on the top right corner.

**(B)** Enrichment of gene ontologies governing development of diverse avian cell types are efficiently recovered by TRIAGE compared to original expression or FH score. Data show enrichment of embryonic development GO terms for developing chicken embryo data (Rehimi et al., 2016) (Fisher's exact test, one-tailed). Genes are ranked by TRIAGE, functional heterogeneity (FH) score, or expression value and binned into rank bins. For the full list of GO terms compared, see **Table S7**.

**(C)** TRIAGE is efficient at identifying developmental regulators compared to using differential H3K27me3 or original expression. Data show functional enrichment of heart-specific GO developmental terms among top 100 genes ranked by TRIAGE, H3K27me3 loss (ΔH3K27me3) or expression value, for cardiomyocyte data (Paige et al., 2012). For the full GO term list, see **Table S8**.

**(D)** TRIAGE prioritized genes are accounted for only in part by analysis of differential H3K27me3 during cell differentiation. Data show a scatter plot for the discordance score and H3K27me3 difference (ΔH3K27me3, between day0 and day14) demonstrating that differential H3K27me3 does not entirely account for genes prioritized by TRIAGE.

**(E)** TRIAGE is sensitive in detecting genes controlling development of diverse cell types in mesendoderm development. Data show enrichment of stage-specific developmental GO terms in TFs ranked by the expression value, TRIAGE or fold change from differential expression (DE) analysis between different clusters defined in a previous study (Friedman et al., 2018). The enrichment analysis (Fisher's exact test, one-tailed) is performed across rank positions (x-axis). Only TFs are included in the analysis. Bar plots show enrichment of other related GO terms at the top 1% rank position. For the full list, see **Table S9**.

**(F)** Brain, heart, and blood RNA-seq samples show enrichment of tissue-specific TFs in genes ranked by TRIAGE but not by original expression or DE fold change. The tissue-specific TFs are defined as those within each graph's specified GO term. The DE ranking is based on the fold change between input sample and other tissue groups shown.

**A** **Rank list of genes from original input expression or TRIAGE analysis of mesendoderm cell types**
Friedman *et al*, Cell Stem Cell, 2018

***Day 2 Cluster 1 Definitive Endoderm***

| Expression rank | Expression gene | TRIAGE rank | Expression rank | TRIAGE gene | TRIAGE rank |
|---|---|---|---|---|---|
| 1 | EEF1A1 | 486 | 55 | SOX17 | 1 |
| 2 | MALAT1 | 1021 | 499 | LINC00261 | 2 |
| 3 | GAPDH | 1022 | 733 | OTX2 | 3 |
| 4 | FTL | 1023 | 1254 | PRAC1 | 4 |
| 5 | FTH1 | 1024 | 238 | GAD1 | 5 |
| 6 | LDHA | 487 | 1297 | TBX3 | 6 |
| 7 | BNIP3 | 249 | 587 | EOMES | 7 |
| 8 | RACK1 | 1025 | 42 | CYP26A1 | 8 |
| 9 | TMSB4X | 130 | 225 | GNAS | 9 |
| 10 | NPM1 | 488 | 2669 | SIX3 | 10 |

● Endoderm related transcription factor, known
● Endoderm regulatory gene, known
● All others, including unknown endoderm genes

☒ Gene rank position based on expression
☒ Gene rank position based on discordance score

***Day 2 Cluster 2 Mesoderm***

| Expression rank | Expression gene | TRIAGE rank | Expression rank | TRIAGE gene | TRIAGE rank |
|---|---|---|---|---|---|
| 1 | EEF1A1 | 492 | 632 | MSX1 | 1 |
| 2 | GAPDH | 1017 | 1052 | PRAC1 | 2 |
| 3 | MALAT1 | 1018 | 859 | OTX2 | 3 |
| 4 | FTL | 1019 | 560 | EOMES | 4 |
| 5 | NPM1 | 493 | 537 | GAD1 | 5 |
| 6 | RACK1 | 1020 | 222 | GNAS | 6 |
| 7 | FTH1 | 1021 | 3229 | GATA4 | 7 |
| 8 | PTMA | 1022 | 1971 | TBX3 | 8 |
| 9 | BNIP3 | 268 | 1097 | CNTFR | 9 |
| 10 | HNRNPA1 | 1023 | 2079 | HAND1 | 10 |

● Mesoderm related transcription factor, known
● All others, including unknown mesoderm genes

☒ Gene rank position based on expression
☒ Gene rank position based on discordance score

***Day 5 Cluster 1 Cardiac Progenitor***

| Expression rank | Expression gene | TRIAGE rank | Expression rank | TRIAGE gene | TRIAGE rank |
|---|---|---|---|---|---|
| 1 | MALAT1* | 930 | 136 | HAND1* | 1 |
| 2 | EEF1A1 | 931 | 1069 | GATA4* | 2 |
| 3 | TPM1* | 53 | 553 | COL2A1 | 3 |
| 4 | MDK | 445 | 1310 | PITX2* | 4 |
| 5 | FTH1 | 932 | 196 | GNAS* | 5 |
| 6 | VIM | 223 | 1581 | NKX2-5* | 6 |
| 7 | KRT18 | 446 | 606 | IGF2* | 7 |
| 8 | FTL | 933 | 2061 | MSX1* | 8 |
| 9 | ACTC1* | 47 | 925 | ISL1* | 9 |
| 10 | RACK1 | 934 | 209 | BMP4* | 10 |

● Cardiac transcription factor, known
● Cardiac regulatory gene, known
● Cardiac structural gene, known
● All others, including unknown cardiac genes
* Indicates genes implicated in cardiac disease

☒ Gene rank position based on expression
☒ Gene rank position based on discordance score

***Day 5 Cluster 3 Endothelial precursor***

| Expression rank | Expression gene | TRIAGE rank | Expression rank | TRIAGE gene | TRIAGE rank |
|---|---|---|---|---|---|
| 1 | MALAT1 | 914 | 1971 | GATA4* | 1 |
| 2 | EEF1A1 | 915 | 1647 | GATA2* | 2 |
| 3 | PTMA | 916 | 1233 | TAL1* | 3 |
| 4 | FTH1 | 917 | 436 | GNAS* | 4 |
| 5 | GAPDH | 918 | 1442 | COL2A1 | 5 |
| 6 | TUBA1B | 919 | 2332 | HAND1* | 6 |
| 7 | MYL6* | 920 | 2946 | NKX2-5* | 7 |
| 8 | RACK1 | 921 | 2984 | GATA3* | 8 |
| 9 | FTL | 922 | 2907 | MSX1* | 9 |
| 10 | MDK | 445 | 1290 | BUB3 | 10 |

● Endothelium related transcription factor, known
● Endothelial regulatory gene, known
● Structural gene, known
● All others, including unknown cardiac genes
* Indicates genes implicated in disease

☒ Gene rank position based on expression
☒ Gene rank position based on discordance score

***Day 15 Cluster 1 Non-contractile cardiac derivative***

| Expression rank | Expression gene | TRIAGE rank | Expression rank | TRIAGE gene | TRIAGE rank |
|---|---|---|---|---|---|
| 1 | MALAT1 | 831 | 255 | HAND1* | 1 |
| 2 | EEF1A1 | 832 | 85 | IGF2* | 2 |
| 3 | FTL | 833 | 730 | PITX2* | 3 |
| 4 | KRT18 | 395 | 2010 | GATA4* | 4 |
| 5 | TMSB4X | 129 | 174 | GNAS* | 5 |
| 6 | FTH1 | 834 | 1422 | COL2A1 | 6 |
| 7 | ACTG1 | 835 | 2298 | SCRG1 | 7 |
| 8 | RACK1 | 836 | 238 | DLK1 | 8 |
| 9 | GAPDH | 837 | 1678 | HAND2* | 9 |
| 10 | TAGLN | 838 | 2522 | WT1* | 10 |

● Cardiac transcription factor, known
● Cardiac regulatory gene, known
● Cardiac structural gene, known
● All others, including unknown cardiac genes
* Indicates genes implicated in cardiac disease

☒ Gene rank position based on expression
☒ Gene rank position based on discordance score

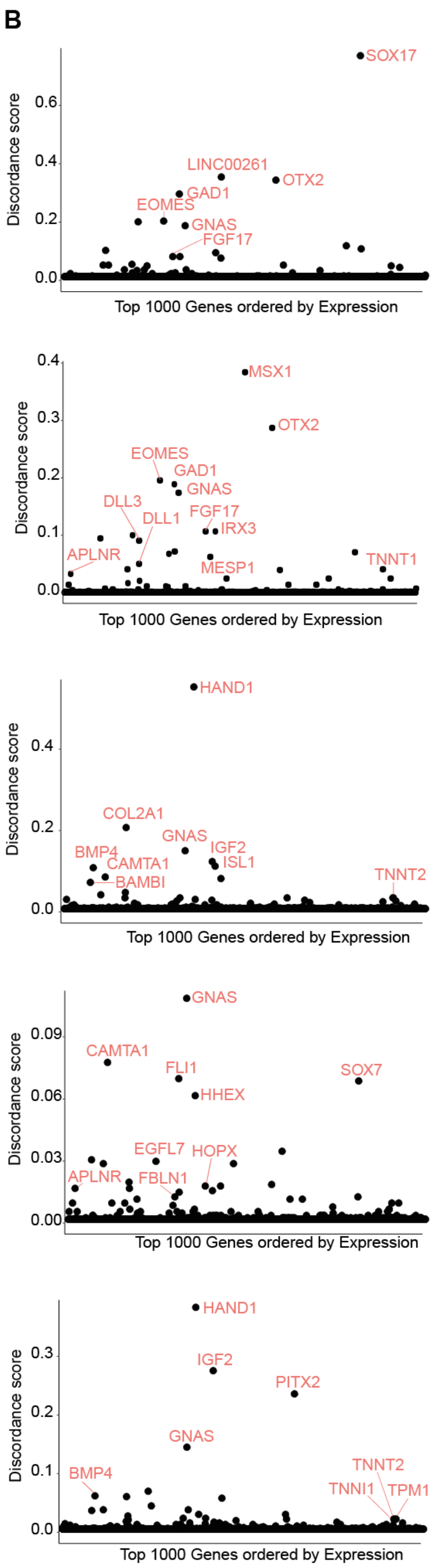


**Figure S6. TRIAGE identifies known cell-type specific regulatory genes across diverse cell populations found during *in vitro* cardiac-directed differentiation (Related to Figure 5).**
**(A-B)** Top 10 genes ranked by the expression value or the discordance score **(A)** and transformation of the transcriptomic expression profile to the discordance score **(B)** from diverse cell populations from scRNA-seq of *in vitro* directed cardiac differentiation during germ layer specification (Day-2), cardiac progenitor specification (Day-5) and cardiomyocyte specification (Day-30).

A Analysis of GTEx data by TRIAGE showing robust enrichment of tissue and cell specific gene ontologies

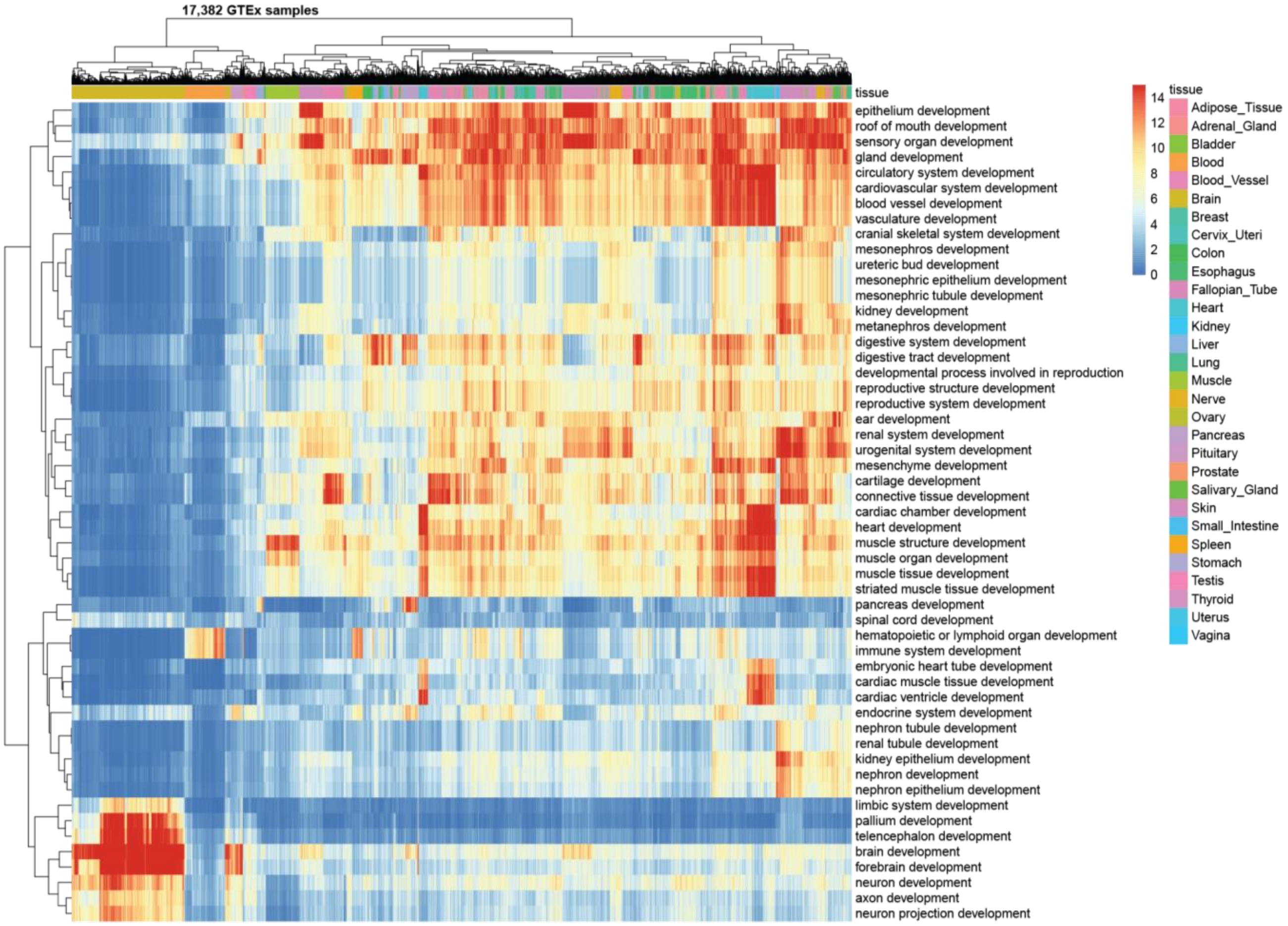


B Analysis of GTEx data using original input gene expression showing poor enrichment of tissue and cell specific gene ontologies

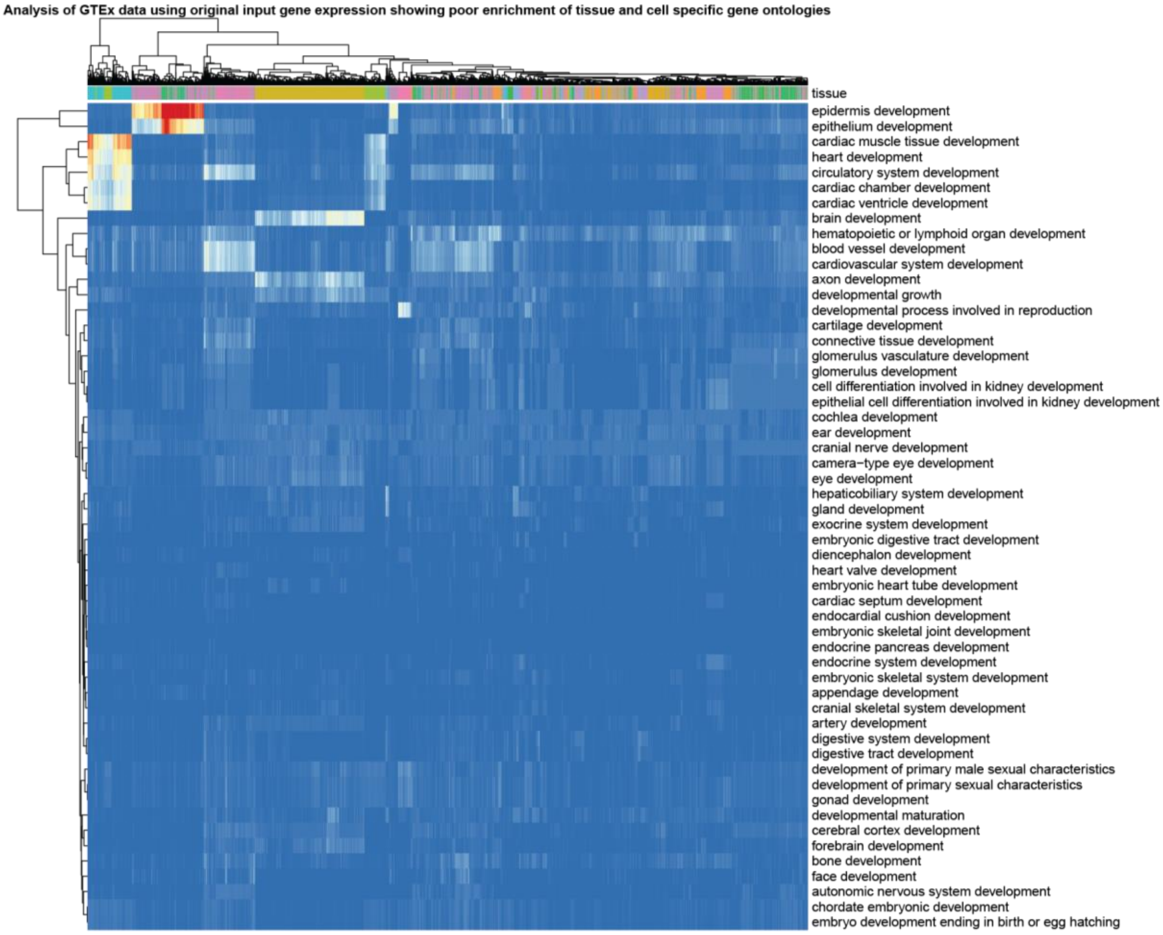

**Figure S7. TRIAGE identifies tissue-specific developmental genes across 17,382 GTEx (v8) samples from diverse tissue and cell types (Related to Figures 4 and 5).**

**(A)** Enrichment of tissue-specific developmental GO terms associated with top 100 genes identified by TRIAGE. For the plot. 53 terms that meet following two conditions are selected: (i) the GO term contains 'development' in the description, and (ii) the GO term is enriched specifically in less than 25% of all GTEx samples (Benamini-Hochberg FDR<$1e^{-15}$, hypergeometric test). Depicted values are -log10 (FDR).

**(B)** Enrichment of the same GO terms among top 100 genes by the expression value. Data are detailed in **Table S11**.

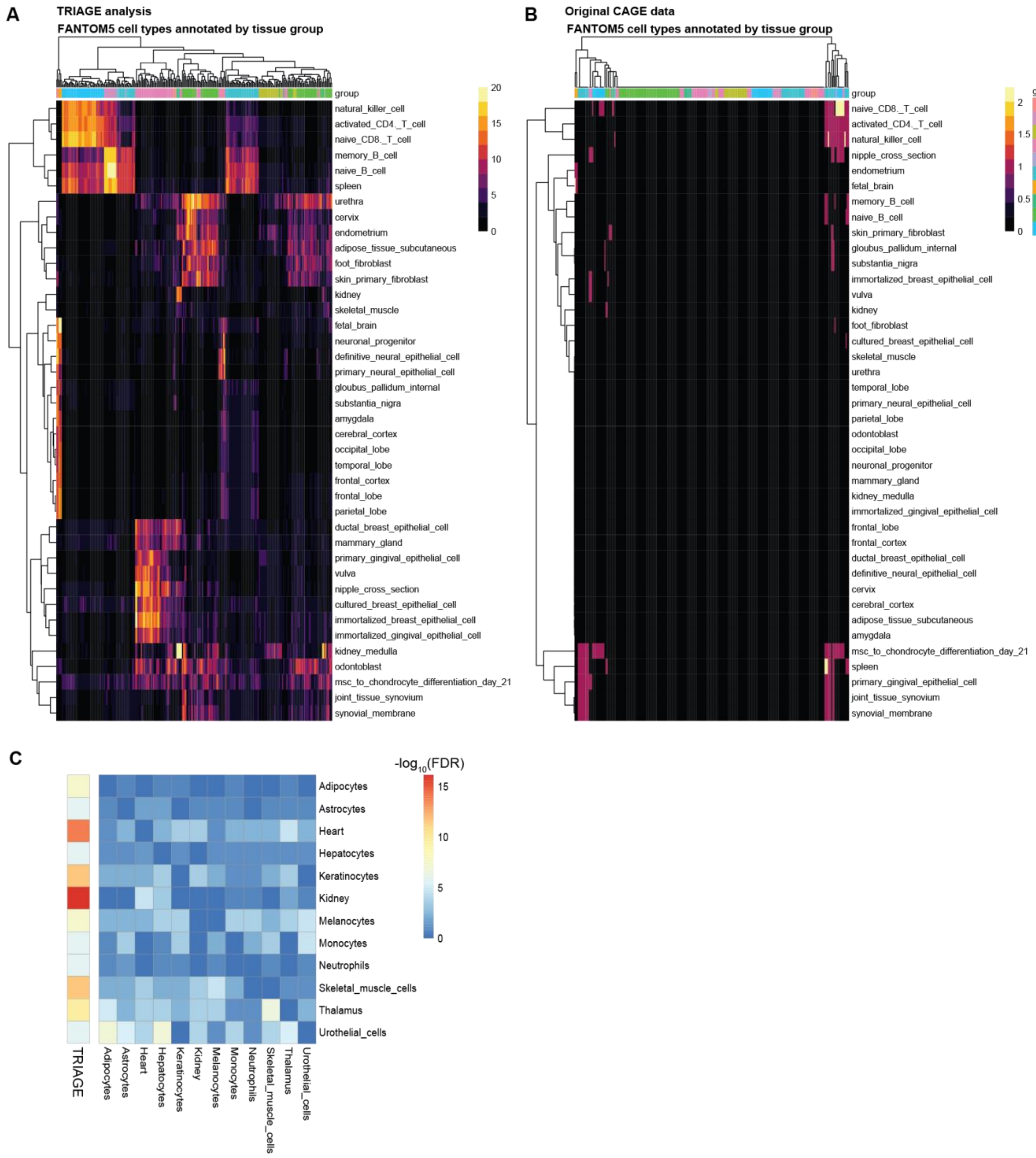


**Figure S8. TRIAGE enriches tissue-specific TFs across 248 FANTOM5 CAGE-seq samples.**
**(A-B)** Tissue-specific TFs are top 20 most tissue-specific TFs defined previously (D'Alessio et al., 2015). Enrichment of the tissue-specific TFs (hypergeometric test) among top by **(A)** TRIAGE or **(B)** expression value. Depicted values are -log10(FDR). Data are detailed in Table **S13**.
**(C)** Analysis of 12 FANTOM5 cell types analyzed by TRIAGE or differential expression shows that TRIAGE consistently and effectively enriches for cell type-specific transcription factors (D'Alessio et al., 2015) among top 100 genes more efficiently than any pairwise comparison using differential expression.

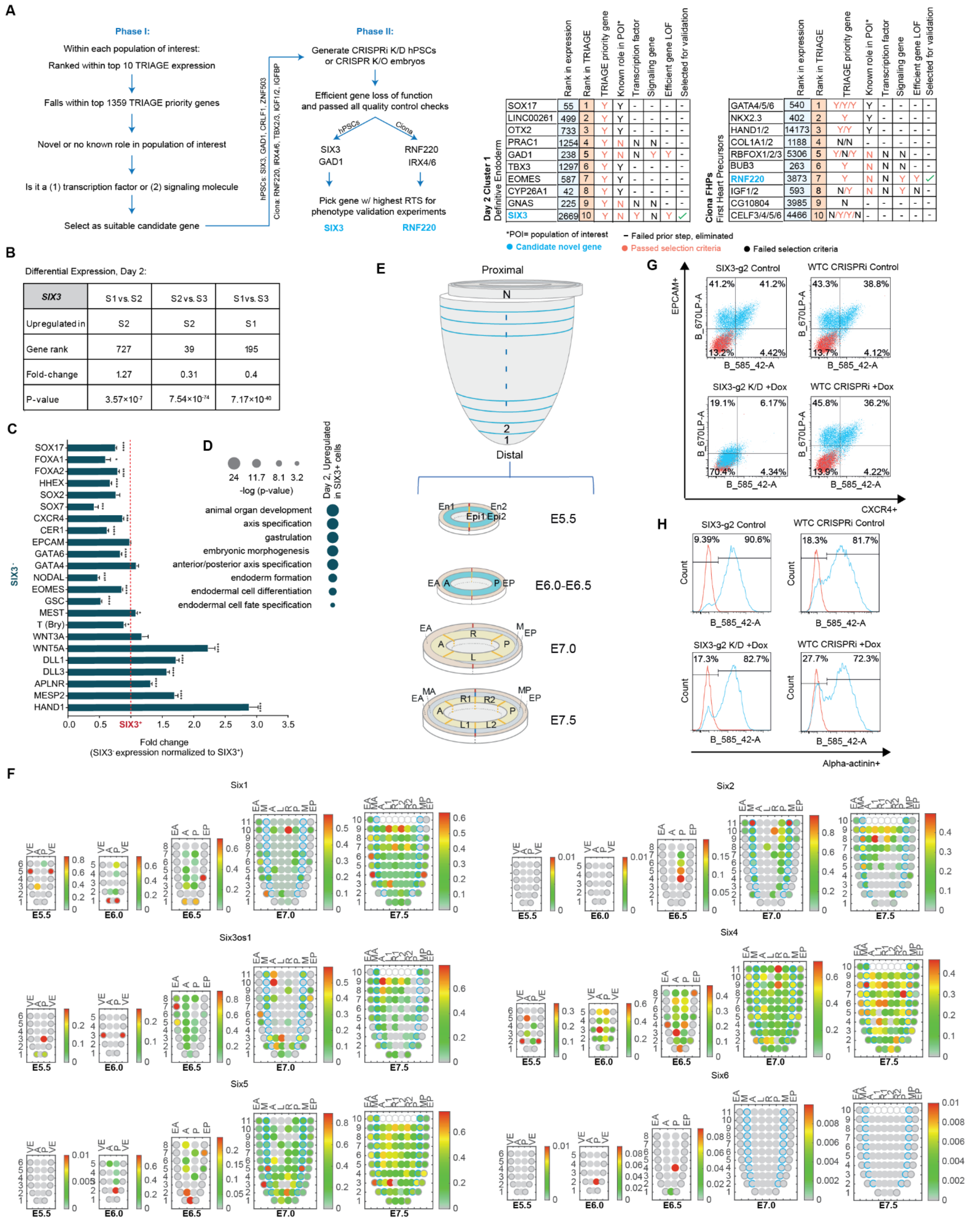

**Figure S9. Biological validation of TRIAGE predictions (Related to Figure 6).**
**(A)** We outline a rubric for gene discovery. **Step 1.** Identify highly ranked TRIAGE genes based on discordance score which would be predicted to play regulatory role. **Step 2.** Is the gene within the top 1359 priority TRIAGE genes to avoid false positives that could be highly ranked based on high abundance gene expression only. **Step 3.** Identify evidence in the literature that the gene as a candidate regulator of the cell type of interest to reinforce the findings from independent studies. **Step 4.** Identify evidence of whether the gene plays a role in diseases associated with mesendoderm

to provide additional evidence of an important role in organismal development and cell differentiation. **Step 5.** Determine if the gene has a functionally validated role in cells where the gene is expressed to ensure no other studies have performed loss of function studies of the candidate gene. **Step 6.** Carefully evaluate the candidate gene biological function as it matches with phenotypic endpoints. **Step 7.** Evaluate and quality control gene loss of function model for functional studies. Schematic shown (Left) provides detailed selection workflow of candidate genes for biological validation. (Right) Detailed criteria used to select SIX3 and RNF220 to be validated as novel regulators of cell identity in hPSCs and *Ciona*, respectively.
**(B)** *SIX3* gene rank using differential expression analysis comparing clusters on Day-2 of scRNA-seq of *in vitro* cardiac directed differentiation shows that SIX3 was not prioritized in this expression-based analysis.
**(C)** Analysis of gene expression comparing *SIX3*$^{+}$ vs *SIX3*$^{-}$ cells on Day 2 of scRNA-seq dataset assessing a panel of germ layer specification genes including markers of endodermal lineages (*SOX17, FOXA2, FOXA1, HHEX, SOX2, SOX7, CXCR4, CER1, GATA6*), mesendoderm (*EPCAM. NODAL, EOMES, GSC*) and mesodermal lineages (*GATA4, T-Bry, WNT3A, WNT5A, DLL1, DLL3, APLNR, MESP2, HAND1*).
**(D)** Gene ontology (GO) analysis on day-2 of differentially expressed genes between *SIX3*$^{+}$/$^{-}$ populations displaying GO terms upregulated in *SIX3*$^{+}$ cells.
**(E)** Sample collection from E5.5-E7.5 embryos for analysis of spatiotemporal transcription. Positions of the cell populations in the embryo: the proximal-distal location in descending numerical order (1 = most distal site, N value of the most proximal section varied by the proximal-distal size of the embryo) and in the transverse plane of the germ layers: endoderm, anterior half (EA) and posterior half (EP); mesoderm, anterior half (MA) and posterior half (MP); epiblast/ectoderm, anterior (A), posterior (P) containing the primitive streak, right (R)-anterior (R1) and posterior (R2), left (L)-anterior (L1) and posterior (L2).
**(F)** Corn plots showing spatial domains of *SIX* family genes expression in the germ layers of E5.5, E6.0, E6.5, E7.0 and E7.5 mouse embryos. The "kernels" in the plot represent the cell populations at different positions in the tissue layers of the embryo (panel **F**). Gradient scale shows the level of gene expression (by RNA-seq transcript reads) in each kernel.
**(G)** Raw FACS plots of EPCAM/CXCR4 analysis for all conditions tested (*n*=12-16 technical replicates per condition from 4-5 experiments).
**(H)** Raw FACS plots of α-actinin analysis for all conditions tested (*n*=6 technical replicates per condition from 3 experiments).

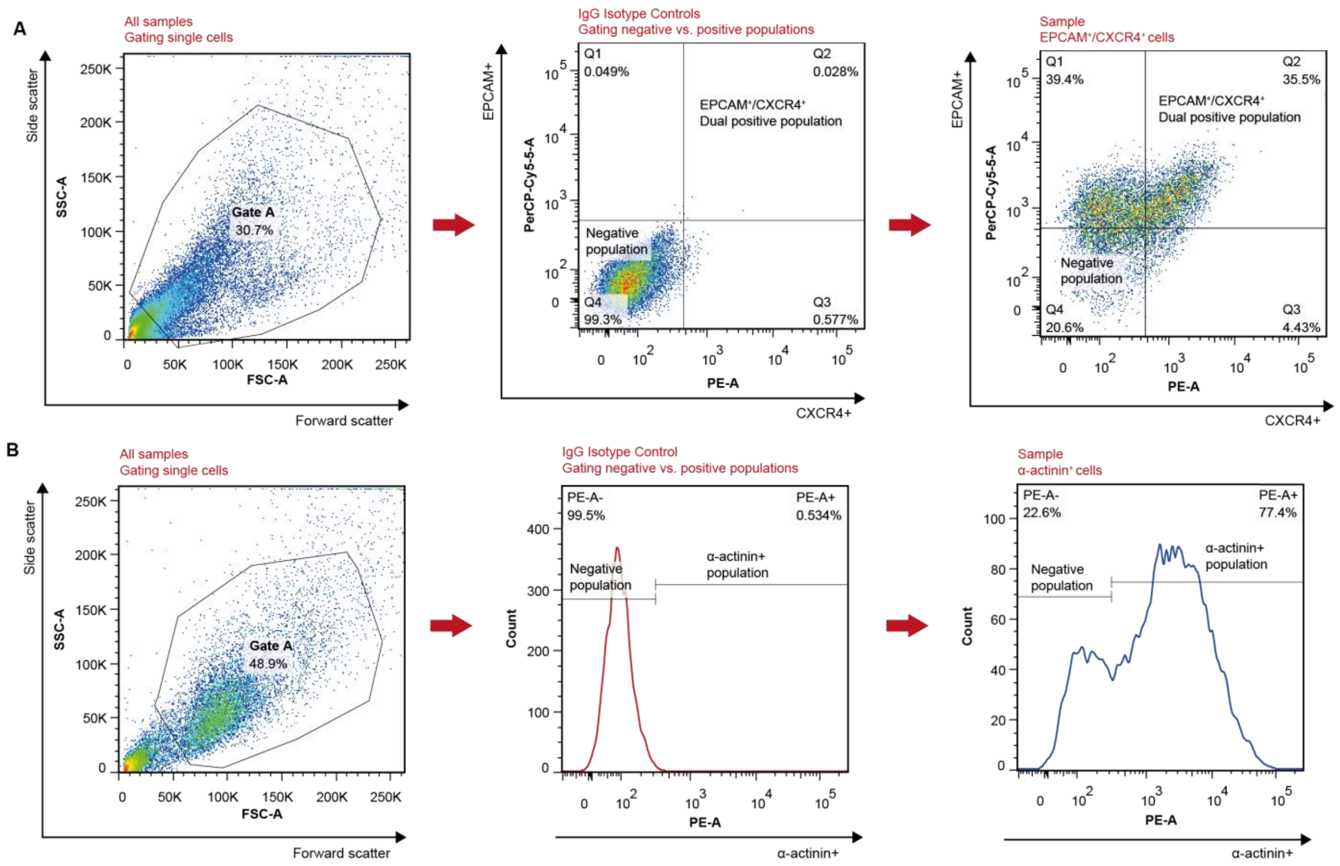


**Figure S10. Flow cytometry gating for analyses of SIX3-KD hPSCs (Related to Figure 6).**

**(A)** For Day-2 samples dual-stained for EPCAM/CXCR4: first, all samples were gated to select singlets, excluding debris and doublets (Gate A= SSC-A vs. FSC-A). Singlets from fluorophore-specific IgG isotype controls (negative control) were used to distinguish between negative and positive populations in two parameter density plots (PerCP-Cy5.5-A vs. PE-A).

**(B)** For Day-15 samples stained for alpha-actinin: first, all samples were gated to select singlets (Gate A= SSC-A vs. FSC-A). Singlets from fluorophore-specific IgG isotype controls were used to distinguish between negative and positive populations in single parameter histograms (Count vs. PE-A).

**Supplementary Tables**

**Table S1.** Descriptions of 111 NIH Roadmap consolidated epigenomes, FANTOM5, human proteomics and inter-species datasets used in the study.

**Table S2.** Lists of 634 variably expressed TFs (VETFs)

**Table S3.** Details of broad H3K27me3 domains from the 111 Roadmap cell types and the repressive tendency score (RTS) table.

**Table S4.** Enrichment of KEGG pathway terms and selected GO biological process terms. Genes were ranked by the RTS and enrichment of a specific KEGG pathway or GO BP term was analyzed using Fisher's exact test (one-sided) across rank positions.

**Table S5.** Lists of ranked genes by different methods (i.e. TRIAGE, FH score, nearest active genes to SEs and H3K4me3) across the 5 tissue types.

**Table S6**. Functional enrichment for top ranked *n* genes (where *n* is the number of active genes nearest to super-enhancers, see Methods), comparing between TRIAGE and SE-based approach across 5 different Roadmap tissue types. Values shown are the false discovery rate by Benjamini-Hochberg method.

**Table S7**. Enrichment of 19 embryonic neural differentiation GO terms across rank positions (columns), comparing the performance between TRIAGE, functional heterogeneity (FH) score and expression value (Rehimi et al., 2016). Values shown are *p*-value at a given rank percentile bin (Fisher's exact test, one-tailed).

**Table S8**. Functional enrichment for top 100 genes ranked by TRIAGE, H3K27me3 loss (between day 0 and definitive cardiomyocyte, day 14) or expression value for definitive cardiomyocyte data (day 14) (Paige et al., 2012) (Fisher's exact test, one-sided).

**Table S9**. Functional enrichment for top 1% TFs ranked by TRIAGE, fold-change from differentially expressed (DE) gene analysis or expression value. For the analysis, only TFs are included.

**Table S10**. Enrichment of GO BP terms (shown as -log10 (Benjamini-Hochberg FDR), hypergeometric test) associated with top 100 genes by discordance score (TRIAGE) or expression value for the Tabula Muris data.

**Table S11**. Enrichment of GO BP developmental terms associated with top 100 genes by discordance score (TRIAGE) or expression value for 17,382 GTEx (v8) transcriptome data. Shown values are average significance (-log10 (Benjamini-Hochberg FDR), hypergeometric test) for the tissue group. Also, the proportion of samples that are enriched (FDR<1e-6) with a given GO term in each tissue group is included.

**Table S12**. Enrichment of GO BP developmental terms (shown as -log10 (Benjamini-Hochberg FDR), hypergeometric test) associated with top 100 genes by discordance score (TRIAGE) or tag density for Roadmap H3K36me3 data.

**Table S13**. Enrichment of tissue-specific TFs from 233 different tissue groups (rows) across 248 FANTOM5 CAGE-seq samples (columns). Top 20 most tissue-specific TFs are used as the positive gene set (D'Alessio et al., 2015). The enrichment compares top 100 genes identified by discordance score (TRIAGE) or expression value for FANTOM5 CAGE values. Depicted values are -log10 (Benjamini-Hochberg FDR, hypergeometric test).

**Table S14**. Enrichment of GO BP developmental terms (shown as -log10 (Benjamini-Hochberg FDR), hypergeometric test) associated with top 100 genes by discordance score (TRIAGE) or expression value for 329 FANTOM5 CAGE-seq samples.

**Table S15**. Enrichment of GO BP developmental terms (shown as -log10 (Benjamini-Hochberg FDR), hypergeometric test) associated with top 100 genes by discordance score (TRIAGE) or expression value for human proteomic data.
